# Data-driven multiscale modeling deciphers MOI-dependent dual antiviral mechanisms of OP7

**DOI:** 10.64898/2026.09.08.750025

**Authors:** D. Rüdiger, P. Opitz, J. Küchler, U. Reichl, S. Y. Kupke

## Abstract

OP7 defective interfering particles are promising antivirals against influenza A virus (IAV), but their antiviral mechanisms are not fully understood. Here, we developed a data-driven multiscale model of IAV and OP7 coinfection calibrated to *in vitro* human lung cell data. The model predicted, and experiments confirmed, a previously unrecognized multiplicity of infection (MOI)-dependent switch in the relative contribution of two complementary antiviral mechanisms of OP7. For low IAV MOI, OP7-mediated interferon signaling induces the antiviral effector MxA, restricting nuclear import of IAV genomes, while replication interference subsequently reinforces complete suppression of virus replication. For high MOI coinfection, the interferon response established is too slow to contribute to antiviral activity, and virus inhibition is mediated almost exclusively by replication interference. The model further predicted a therapeutic efficacy of OP7 up to 24 h post infection, which was confirmed in subsequent experiments. Following extension, it also reproduced the experimentally defined prophylactic protection for up to 7 days. Altogether, this experimentally validated coinfection model provides a quantitative framework for understanding and rationally optimizing prospective OP7-based antiviral therapies.

## Introduction

Viral infections have resurfaced as a major topic of concern in recent years due to the short- and long-term threats they pose to human health and the global economy (1). Vaccination remains the most effective and efficient countermeasure against known diseases, but might lag behind significantly when new viruses or potent mutants emerge. Antiviral agents can bridge this gap, but often target specific pathogens constraining their suitability against diverse respiratory infections (2). Additionally, their efficacy might decline due to emerging antiviral resistances (3, 4). Broad-spectrum antivirals that inhibit various virus species could provide an additional safeguard in case of pandemic outbreaks or bioweapon deployment (5). Currently, over 100 broad-spectrum antivirals are in development, but there does not seem to be a blockbuster similar to penicillin in sight (2). As the recent COVID-19 pandemic has clearly demonstrated, an expansion of our current treatment catalogue for virus infections is of utmost importance. Influenza A virus (IAV)-derived defective interfering particles (DIPs) have been proposed as promising broad-spectrum antiviral agents based on their capability to suppress IAV and multiple unrelated virus species, e.g., influenza B virus, yellow fever virus, respiratory syncytial virus, Zika virus, and SARS-CoV2 (6–8).

DIPs are replication-incompetent derivates of their corresponding standard virus (STV) (9–17). They lack critical functions required to propagate on their own but can replicate during coinfection with their STV. For conventional IAV DIPs, this defect is caused by a large internal deletion in (at least) one of the eight viral genome segments. In a coinfection scenario, the STV provides the missing functions and, due to their reduced length, DIP viral genomic RNAs (vRNAs) replicate faster accumulating to higher levels than STV vRNAs. This causes the depletion of viral resources and, thereby, suppress the propagation of the STV, a process referred to as “replication interference”. (10, 12, 18, 19). Additionally, DIPs were shown to stimulate antiviral innate immune responses, in particular, the interferon (IFN) system, leading to the induction of an antiviral state in DIP-infected cells (20–22). Due to these properties, DIPs of various virus species were studied for antiviral therapy (23–27). The antiviral capabilities and a safe profile upon administration of DIPs have been demonstrated in various animal models (28–35). However, it remains a matter of debate which antiviral mechanism of DIPs, replication interference or IFN-mediated activity, plays the dominant role in suppressing virus infections (8, 23, 34, 36–39).

OP7, an alternative type of IAV DIP, was discovered previously by our group and showed even greater antiviral effects than conventional DIPs in cell culture and animal studies (7, 8, 28, 29, 40–42). OP7 does not contain a large internal deletion but carries several point mutations in the sequence of its segment 7 (S7) vRNA. S7 of the STV (S7-STV) encodes for the structural and regulatory matrix protein 1 (M1) that is crucial for viral uncoating, nuclear export and progeny virion formation. Despite its mutations, S7 of OP7 (S7-OP7) vRNA is still capable to induce mRNA transcription and the synthesis of M1-OP7 protein, whose functionality is likely affected by the mutations (43). Previously, we explored how OP7 induces the observed replication interference. Some point mutations form a so-called “superpromoter”, which strongly enhances the replication of S7-OP7 vRNA leading to the depletion of viral resources and inhibiting STV propagation (44, 45). Furthermore, the produced M1-OP7 is likely unable to bind vRNAs in the nucleus to initiate their export, which would explain OP7’s defect in virus replication (43). The antiviral efficacy of OP7 was initially tested in cell culture and, subsequently, in animal studies showing that intranasal administration is well-tolerated and can rescue mice infected with lethal IAV doses (8, 28, 40, 41).

Mathematical modeling has been used extensively to describe and predict influenza virus dynamics. The majority of studies focused on population dynamics, be it human populations (46–48) or cell populations (49–54). Other models studied the intracellular processes during infection in depth (55, 56). In addition, we have developed models describing the interaction between STV and DIPs on the intracellular level (18) and on the cell population level (57). Recently, we developed an intracellular model of STV/OP7 coinfection to uncover the effects of the point mutations in S7-OP7 vRNA (43). While this model closely captured single-cycle coinfections, it cannot describe the gradual infection of a cell population starting from low STV multiplicities of infection (MOIs). However, low MOI scenarios are the majority of infection events occurring in nature via airborne transmission (39). Thus, for the description of such scenarios, a multiscale approach covering the intracellular as well as the cell population level has to be employed (58–60). Additionally, the intracellular model of STV/OP7 coinfection did not incorporate the effects of the IFN response as it was calibrated to MDCK cells, which are of canine origin and whose IFN-induced effector proteins do not show activity against human IAVs (61, 62).

In this study, we developed a data-driven multiscale model of STV/OP7 coinfection in IFN-competent human lung cells, calibrated using experimental *in vitro* data that cover a wide range of infection conditions (63). We used this model to investigate how OP7 suppresses IAV replication, to predict therapeutic and prophylactic treatment windows, and to dissect the respective contributions of IFN-mediated inhibition and replication interference to antiviral activity. By integrating quantitative modeling with experimental validation, this work provides mechanistic insight into OP7 antiviral activity and establishes a framework for the rational optimization of OP7-based treatment strategies.

## Results

### A multiscale STV/OP7 coinfection model accurately reproduces virus and IFN dynamics across diverse infection conditions *in vitro*

To investigate the interplay between OP7’s replication interference and antiviral IFN induction, we developed a data-driven multiscale model of STV/OP7 coinfection calibrated with *in vitro* human lung cell data. The framework presented in this study is based on a previously published model describing intracellular coinfection dynamics (43). This model covers the viral life cycle steps in detail, including virus entry, nuclear import, synthesis of viral mRNAs and the vRNA segments, their nuclear export, and progeny virion formation. To cover coinfection dynamics in highly different infection conditions, i.e., combinations of low and high STV MOIs and OP7 doses, and to evaluate the protective effect of OP7 treatment, we extended this model by implementing the cell population level (**Figure 1**). Therefore, we describe the spread of viruses from cell to cell as well as cell growth, (co)infection, apoptosis and lysis. Additionally, we introduced major components of antiviral IFN signaling triggered after infection.

**Figure 1:**
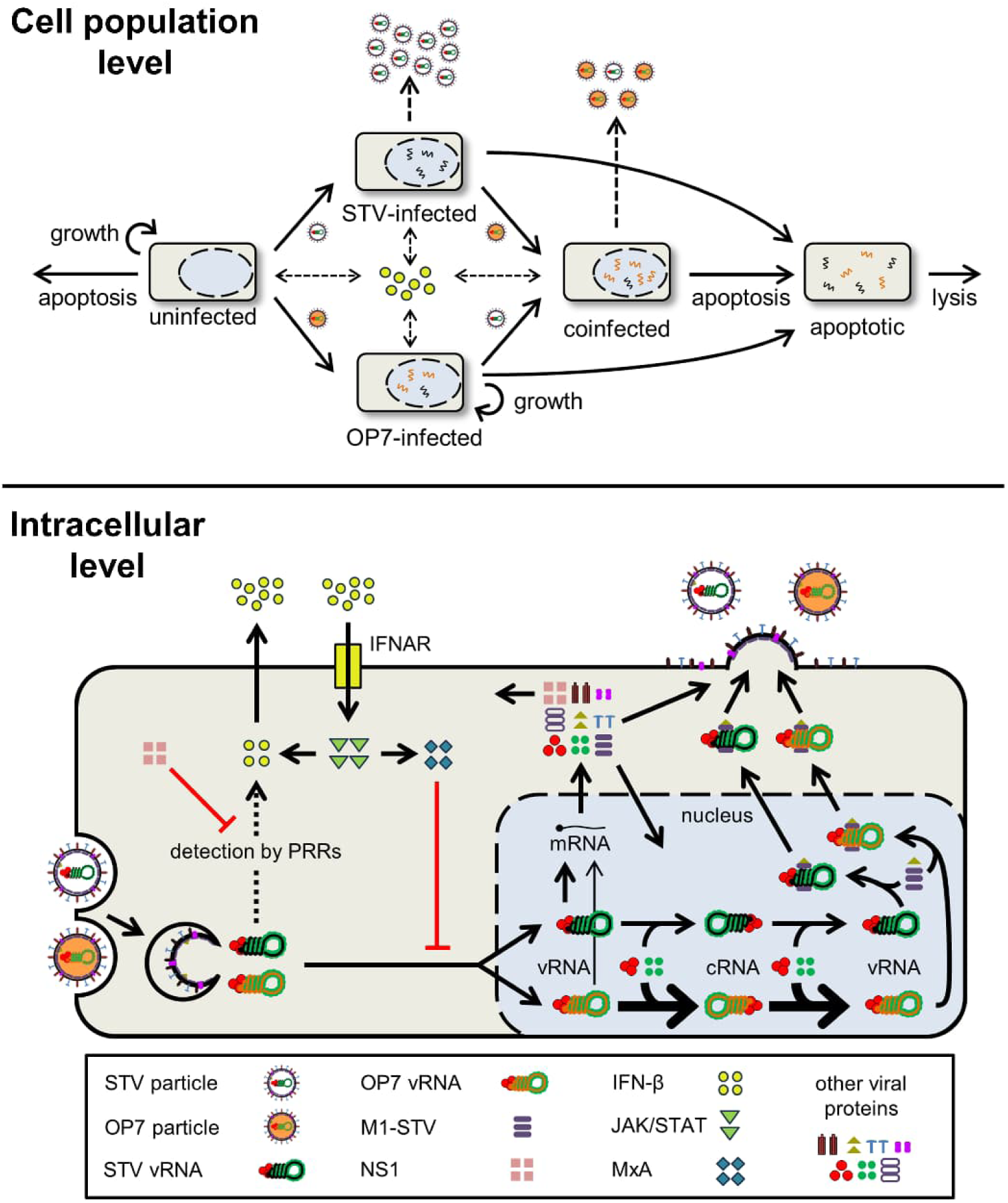
Multiscale model linking intracellular virus replication, IFN signaling and infection spread during STV/OP7 co-infection. (Top) Growth, infection, apoptosis and lysis of uninfected, STV-infected, OP7-infected and coinfected cells are described on the population level. STV particles are released from STV-infected and coinfected cells, OP7 particles are only released from coinfected cells. Both particles are degraded over time. Uninfected cells maintain a baseline level of IFN-β release, while infected cells release increased levels of IFN-β. (Bottom) The intracellular model, which is based on a previous model of OP7 replication (43), is used to simulate virus entry, nuclear import, viral replication and protein synthesis, the IFN response, nuclear export and progeny virion release for both STV-infected and coinfected cells. IFN-β is synthesized and secreted following the detection of incoming vRNAs by PRRs. Then, secreted IFN-β binds to cognate IFNAR receptors on the surface of cells, which initiates the JAK/STAT signaling pathway inducing MxA synthesis. MxA is an antiviral protein whose exact mechanism of action has not been defined, yet. Following model selection, we implemented an inhibition of the nuclear import of vRNAs as the function of MxA. To counteract antiviral IFN signaling, the viral NS1 protein serves as an IFN antagonist that prevents recognition of vRNAs by PRRs.

To describe the IFN system, we focused on the cytokine IFN-β, its cognate cell surface interferon-α/β receptor (IFNAR), the JAK-STAT (Janus kinase, signal transducer and activator of transcription proteins) downstream signaling cascade and the most relevant IFN-stimulated gene (ISGs)-encoded protein, i.e., human myxovirus resistance protein 1 (MxA). During IAV infection, incoming vRNA is detected by pattern recognition receptors (PRRs), e.g., retinoic acid-inducible gene I (RIG-I) and protein kinase R (PKR) (20, 64). The PRRs initiate a signal cascade leading to the expression of IFN-β. It is then secreted to the extracellular space and binds to IFNAR on the surface of other cells (paracrine signaling) and the releasing cell itself (autocrine signaling). These receptors recruit JAK-STAT complexes which initiate the expression of ISGs. While a large variety of ISGs is expressed, the *MX1* gene encoding for MxA is assumed to be a key factor inhibiting IAV (65, 66). However, the exact mode of action of MxA against IAV is still subject of debate (66). After testing the different hypotheses regarding the mode of action of MxA (**Figure 4**), we implemented an inhibition of nuclear import of incoming vRNAs in the model, which will be elaborated on later in this study. Additionally, JAK-STAT signaling induces an increased expression of IFN-α/β genes strengthening the IFN response in reaction to the infection even further (positive feedback loop). Lastly, we introduced the viral NS1 protein, an antagonist of the IFN response which can prevent recognition of vRNAs by PRRs (67, 68).

To calibrate the multiscale model of STV/OP7 coinfection, we used the experimental data published recently by Opitz *et al.* (63). The authors studied the infection dynamics of STV/OP7 coinfection in Calu-3 cells, which are human lung cells capable of inducing an effective IFN response against human IAV infection. This data set includes dynamics of different intracellular vRNAs and viral mRNAs, including S7-STV and S7-OP7, virus titers, viral NS1 protein levels, and the expression of IFN-β and MxA (**Figure 2, Figure S1**). Infections using different initial conditions, i.e., combinations of STV MOIs of 0, 10^-3^, 3, and 30 with no, low and high doses of OP7, were carried out to cover a wide range of scenarios. Overall, the experimental data provided in Opitz *et al.* show that high OP7 doses can completely prevent STV propagation for low MOIs and significantly reduce infectious virus titers for high MOIs (**Figure 2S-X**). Additionally, the (co)infection with OP7 induces a stronger IFN response than STV-only infections (**Figure 2M-R**). This can likely by attributed to the replication interference mediated by OP7 suppressing the accumulation of STV vRNA, which includes S8 encoding for the viral IFN antagonist NS1 in coinfections (63). Lower levels of NS1 would lead to less inhibition of the recognition of vRNAs by PRRs and, in turn, to an increased IFN response.

**Figure 2:**
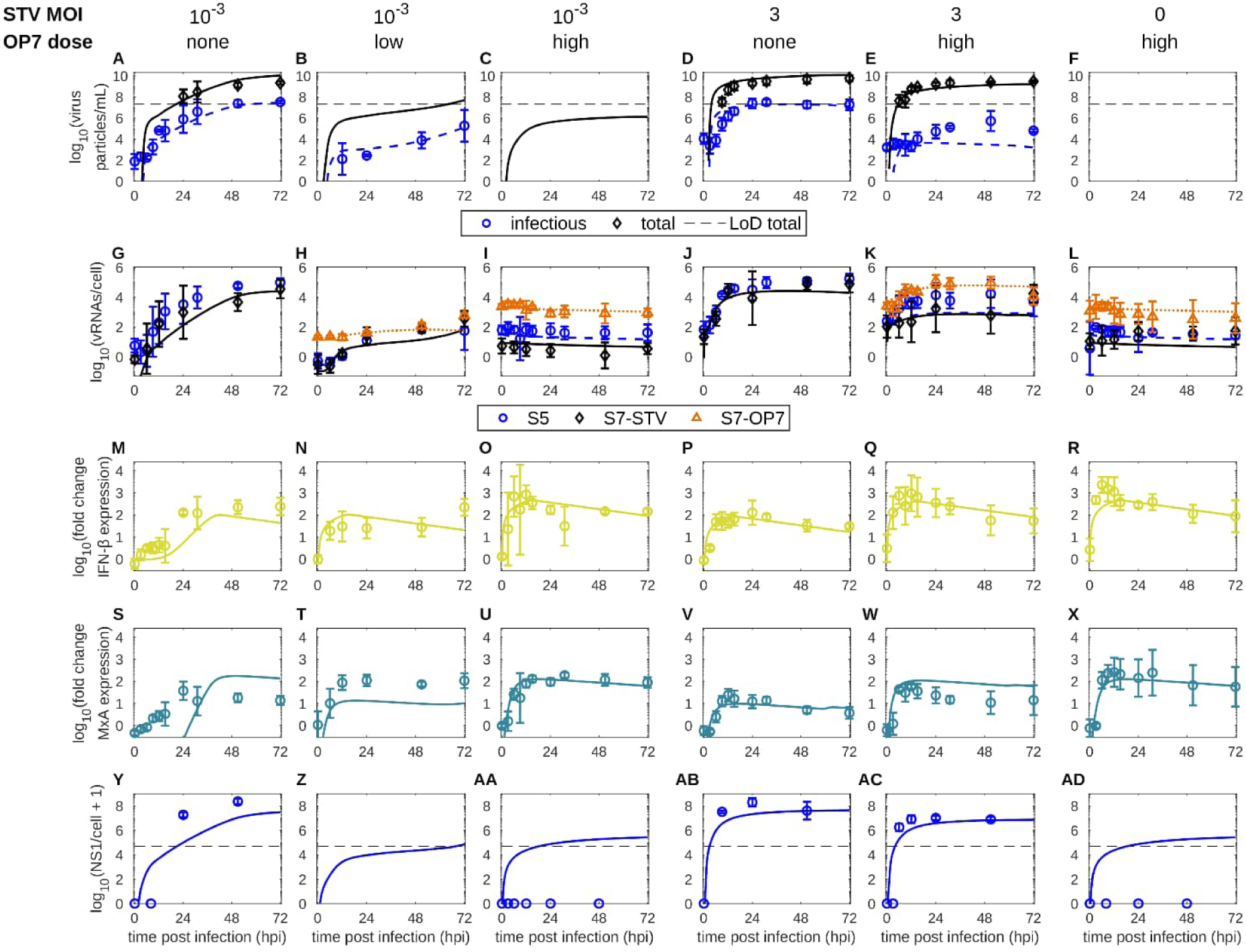
Model simulations accurately reproduce intracellular and extracellular virus dynamics and IFN induction across infection conditions. Simulations of the STV/OP7 coinfection model, fitted to experimental data from infections of human lung cells (63). Cells were infected using combinations of different STV MOIs (influenza A/PR/8/34, H1N1) and OP7 doses. (A-F) Extracellular virus titers, (G-L) cell-specific vRNA levels, (M-R) fold change of IFN-β and (S-X) MxA expression, and (Y-AD) cell-specific viral NS1 protein levels. The black dashed lines indicate the limit of detection (LoD) for (A-F) total virus particles/mL and (Y-AD) viral NS1 protein levels. Error bars show the standard deviation of three independent experiments. See also **Figure S1**.

As an initial step, we used a coinfection model that did not contain the IFN system and calibrated it to the experimental data. It could describe most infection conditions; however, it showed large deviations for a coinfection at STV MOI of 10^-3^ and a low OP7 dose (**Figure S2**). To represent the observed infection dynamics completely, the implementation of the IFN system was necessary. Furthermore, we extended the model by adding an inhibition of the nuclear export of vRNAs in OP7-only infected cells to enable the description of the effect of prophylactic OP7 treatment (**Figure 3C-D**). In the following, we examine these changes individually to show the impact of their inclusion.

**Figure 3:**
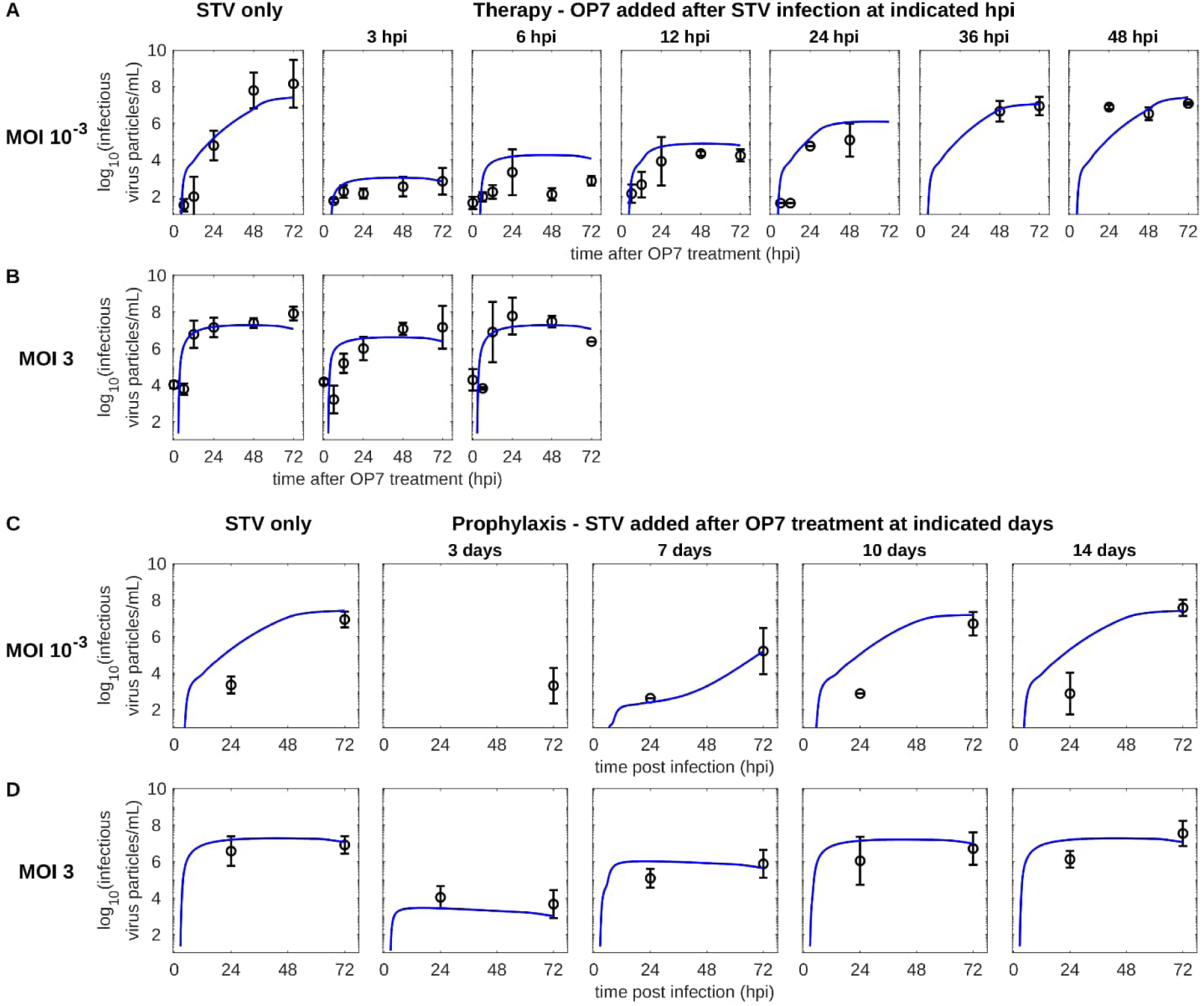
Prediction of the therapeutic antiviral effect and model refinement to explain the prophylactic effect of OP7 application. (A-B) Model prediction, followed by experimental validation, of the therapeutic effect of a high OP7 dose added after STV infection using MOIs of 10^-3^ and 3. (C-D) Experimental outcome of the prophylactic effect of a high OP7 dose added before STV infection using MOIs of 10^-3^ and 3. To reproduce the data, an inhibited export of vRNAs in cells infected only by OP7 was implemented (**Figure S3**). Error bars represent the standard deviation of three independent experiments.

After calibrating the key infection- and IFN-related parameters of the final model (**Figure 1**, **S1 Table**), our simulations captured the complex interdependencies of the virus life cycle and the IFN system for all infection conditions closely. The dynamics of virus titers, vRNA and viral mRNA levels, the change in IFN-β and MxA expression, and viral NS1 protein levels are well represented (**Figure 2, Figure S1**). For STV MOI 3 combined with a high OP7 dose, infectious virus titers are slightly underrepresented in model simulations after 12 hours post infection (hpi) (**Figure 2E**). Furthermore, MxA levels are overestimated in simulations for a STV-only infection with MOI 10^-3^ (**Figure 2S**) and are underestimated in simulations for STV MOI 10^-3^ combined with a low OP7 dose (**Figure 2T**). As the model was calibrated to a wide range of infection conditions at the same time, a few discrepancies could not be avoided, but the general dynamics are represented well.

Overall, the model closely captures virus dynamics and IFN induction across a broad range of infection conditions, providing a quantitative framework for subsequent mechanistic analyses.

### Predictive modeling of therapeutic efficacy and model refinement for reproduction of prophylactic protection

We next assessed whether the calibrated model could predict the therapeutic and prophylactic application windows of OP7. While a predecessor model accurately predicted therapeutic efficacy, refinement was required to explain the experimentally observed long-term prophylactic protection.

First, we examined the therapeutic effect of OP7 on STV infections. We simulated different time points of OP7 treatment after a STV infection at MOI 10^-3^ and 3. Model simulations suggest that for low STV MOIs, a treatment with high OP7 doses induces a strong reduction of infectious virus titers until 12 hpi (**Figure 3A**). For OP7 treatment at 24 hpi, this effect slowly fades and for an addition at 36 hpi no inhibition of infectious titers can be detected anymore. In high MOI conditions, large titer reductions cannot be achieved when OP7 is provided after STV infection (**Figure 3B**). To challenge our prediction, Opitz *et al.* subsequently performed corresponding coinfection experiments determining the therapeutic capabilities of OP7 (63) and we compared them to our simulations. The experimental results were closely predicted by our model (**Figure 3A-B**). Differences in the infectious titer can be observed for OP7 treatment after 6 hpi for a STV MOI of 10^-3^, but all other time points are predicted well.

Similar simulations were performed for the prophylactic application, i.e., an infection with STV after treating cells with a high dose of OP7. Here, a predecessor model initially predicted that virus titers can be reduced by about one log when a STV infection at MOI 10^-3^ occurred 3 days after treatment with OP7, but not thereafter (**Figure S3A**). For a STV MOI of 3, virus titers are barely reduced following prophylactic OP7 treatment (**Figure S3B**). Again, we compared our prediction with experiments Opitz *et al*. performed thereafter (63). However, we observed large differences between the model prediction and the data (**Figure S3A-B**). In the experimental results for both STV MOIs, infectious virus titers were strongly reduced for infections after 3 days and a reduction of one log could still be detected for infections after 7 days following OP7 treatment (**Figure 3C-D**). Thus, our predecessor model appeared to underestimate the antiviral effect of prophylactic OP7 treatment (**Figure S3**).

In order to describe the experimental data, we adjusted our model with respect to the availability of nuclear S7-OP7 vRNA, which was exported too quickly in the predecessor model, leading to diminished antiviral activity following prophylactic OP7 treatment (**Figure S3A-B**). Thus, we introduced an inhibited nuclear export for OP7-only infections (**Figure S3D, Equations (5)-(6)**). This hypothesis is based on a previous study on OP7, suggesting that the export of nuclear vRNAs is inhibited in OP7 infections (40). With this change, the final model can closely describe the prophylactic effect of OP7 for a STV MOI of 3 (**Figure 3C-D**). For a STV MOI of 10^-3^, the reduction of virus titers is well represented for infections at 7 days after OP7 treatment and thereafter. For infections after 3 days, our model simulates a complete inhibition of virus titers, slightly overestimating the prophylactic effect of OP7.

Overall, the calibrated model accurately predicted therapeutic efficacy up to 24 hpi. Following extension, it also reproduced the experimentally observed prophylactic protection for up to 7 days.

### Model selection suggests an inhibition of nuclear import of viral genomes by the IFN-induced MxA

To incorporate IFN-mediated antiviral activity into the final model (**Figure 1**), we evaluated competing mechanistic hypotheses for the antiviral action of MxA, which is the most important ISG-encoded protein for IAV infection (65, 66). Some previous studies suggested that MxA forms dimers that can bind to viral nucleoprotein (NP), which impedes virus replication (69–72). Another set of studies proposed that the inhibition occurs during the nuclear import of vRNAs (73, 74). Lastly, Haller *et al.* suggested that both effects occur simultaneously (66). We implemented these hypotheses into our model individually and estimated the corresponding model parameters (**Table S2**) to the experimental data (**Figure 4**). Here, we focus on coinfections using a STV MOI of 10^-3^ and a low OP7 dose. In this condition, we observed the largest deviation of model simulations and experimental data when fitting a predecessor model lacking an IFN system (**Figure S2**). Thus, we presumed that the inclusion of the IFN system would have the largest impact in this specific setting enabling an assessment of the different hypotheses.

**Figure 4:**
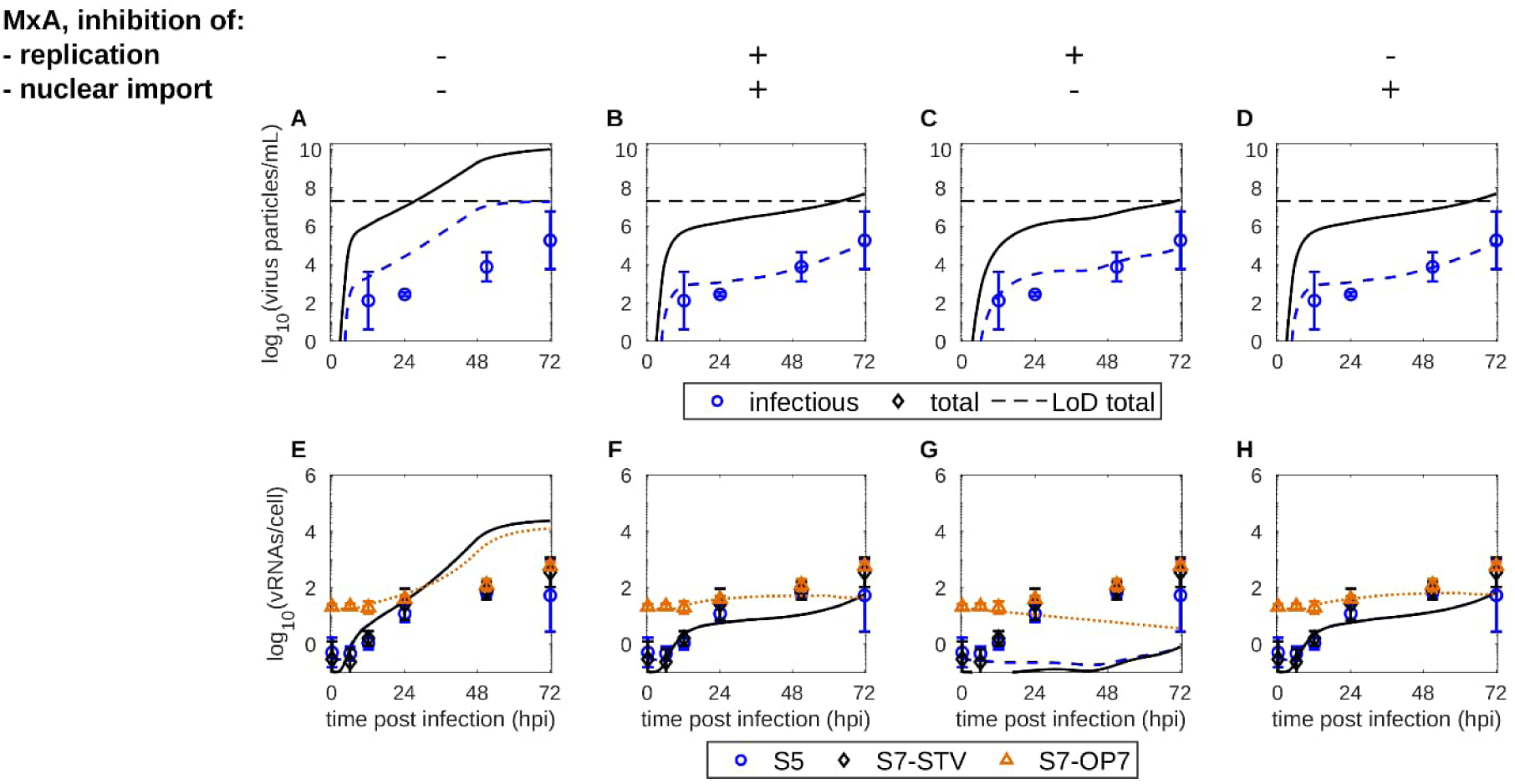
Model comparison identifies inhibition of nuclear import of genomic vRNAs as the most likely mode of MxA action. Simulated virus titers and vRNA levels for a model describing an infection with STV MOI 10^-3^ and a low OP7 dose. The model considers (A+E) the absence of an IFN system, (B+F) the inhibition of virus replication and nuclear import of vRNAs by MxA, (C+G) the inhibition of virus replication by MxA, and (D+H) the inhibition of nuclear import of vRNAs by MxA. The black dashed lines indicate the LoD for total virus particles/mL. Error bars represent the standard deviation of three independent experiments.

The predecessor model without an IFN system could not represent the virus dynamics, which are represented by vRNA levels and virus titers (**Figure 4A+E**). Next, we implemented both mechanisms, which enabled a close description of the virus dynamics (**Figure 4B+F**). To investigate if each mechanism alone was sufficient to explain the observed dynamics, we simulated them separately. Both mechanisms enabled the description of virus titers (**Figure 4C-D**), however, they had significantly different effects on the simulated vRNA levels (**Figure 4G-H**). More specifically, introducing a suppression of virus replication by MxA reduces overall vRNA production significantly, which is not represented in the experimental data (**Figure 4C+G**, **Figure S4**). An inhibition of the nuclear import of vRNAs alone leads to a close description of virus titers and vRNA levels (**Figure 4D+H**). Thus, our model simulations clearly favor the hypothesis that MxA interferes with the nuclear import of vRNA, which is also supported by the Akaike information criterion (**S2 Table**).

In summary, model comparison supports inhibition of nuclear import of incoming genomic vRNAs, rather than viral replication, as the dominant antiviral mechanism of the IFN-induced MxA protein.

### The contribution of IFN-mediated antiviral activity depends on the STV MOI

While the inclusion of the IFN response was required to describe coinfections at STV MOI 10^-3^ and a low OP7 dose (**Figure 4**), coinfections at high STV MOI could already be closely described by a predecessor model lacking an IFN system (**Figure S2**). Thus, we hypothesized that IFN-meditated antiviral activity of OP7 mostly impacts low STV MOI coinfection scenarios. To test this hypothesis, we simulated coinfections using low STV MOI with a low OP7 dose (low MOI coinfection) as well as high STV MOI with a high OP7 dose (high MOI coinfection). Additionally, we tested simulations in which we removed individual sources of the antiviral effect of OP7, i.e., the IFN system and the replication interference of S7-OP7 vRNAs. This replication interference originates from the enhanced replication of S7-OP7 vRNA caused by its “superpromoter”, which leads to the depletion of viral resources and, consequently, the inhibition of STV replication (43). The results of these coinfection simulations are compared to STV-only infections in **Figure 5**.

**Figure 5:**
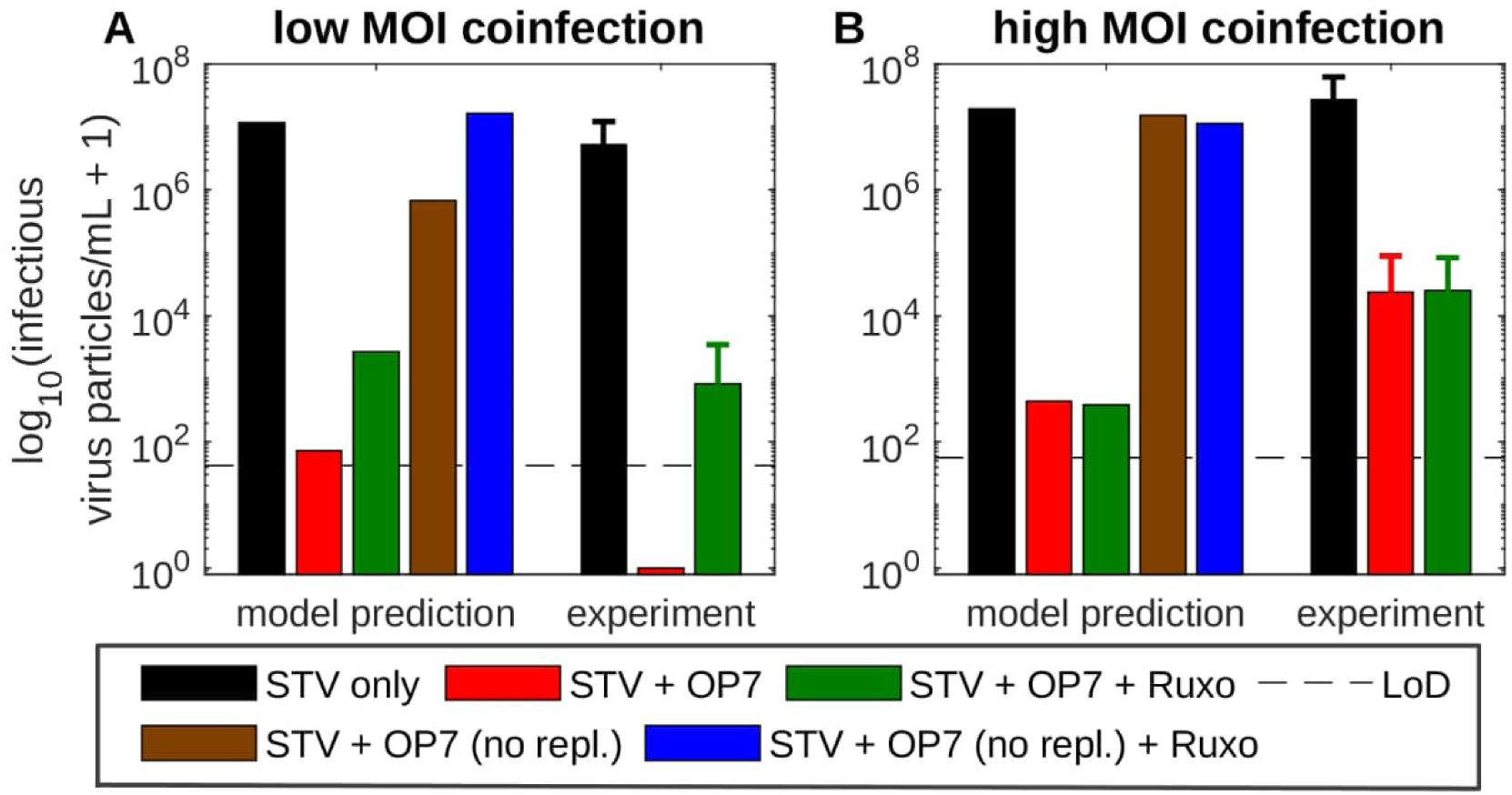
Experimental validation of the predicted MOI dependence of IFN-mediated antiviral activity of OP7. Model simulations and experimental results showing infectious virus titers at 48 hpi resulting from STV-only infection and for different coinfection scenarios in (A) a low MOI, low OP7 dose and (B) a high MOI, high OP7 dose setting. The scenarios include regular coinfections, a coinfection where the IFN response is inhibited by ruxolitnib (Ruxo), a coinfection with a hypothetical OP7 exerting no replication interference (no repl.), and a combination of the latter two effects. Dashed lines represent the LoD for infectious virus particles/mL. Error bars represent the standard deviation of three independent experiments.

The addition of an OP7, which has the full ability to inhibit virus replication, strongly reduced simulated infectious titers (red bars) compared to STV-only infections (black bars) for low and high MOI coinfections (**Figure 5**). Next, model simulations predict that the removal of the IFN system (green bars) only leads to a reduced inhibition of virus release in a low MOI coinfection, but does not affect OP7 antiviral activity in a high MOI coinfection, suggesting that the impact of the IFN response depends on the MOI. On the other hand, simulations with an intact IFN system and a hypothetical OP7 exerting no replication interference (no repl., brown bars) lead to a pronounced reduction of virus titers in a low MOI coinfection and to no inhibition for a high MOI coinfection. This indicates that both mechanisms affect low MOI coinfections. In addition, these simulations suggest that IFN-induced antiviral activity does not contribute to OP7 antiviral activity in high MOI coinfections, where replication interference appears to dominate inhibition instead. Lastly, virus titers reach levels similar to a STV-only infection in low and high MOI coinfections when neither the IFN system nor a replication of S7-OP7 vRNA is present (blue bars).

To test our model prediction, we subsequently performed an infection experiment using ruxolitnib (Ruxo) (63). Ruxo is a JAK inhibitor that halts the activation of the antiviral signaling pathway induced by the IFNAR. Thus, the addition of Ruxo prevents the synthesis of ISG-encoded proteins like MxA, despite the presence of IFNs, which essentially renders IFN-mediated antiviral activity non-functional. In coinfection experiments with Ruxo, a decreased inhibition in a low MOI coinfection, but no impact in a high MOI coinfection, was observed confirming previous model simulations (**Figure 5**).

Taken together, modeling and perturbation experiments demonstrate that IFN-mediated antiviral activity contributes to the antiviral effect of OP7 under low STV MOI coinfection but not at high MOI.

### Replication interference dominates antiviral activity at high STV MOI coinfection as IFN induction is too slow

To decipher the observed MOI dependence of IFN-mediated antiviral activity, we next investigated the temporal dynamics of (co)infection and IFN induction (**Figure 6A-H**). We further evaluated the nuclear import of vRNAs, which is affected by MxA (**Figure 6I-L**). Additionally, we calculated the percentage of viral NP used for the formation of the different vRNAs (**Figure 6M-R**) to examine how much of it is captured by S7-OP7 vRNA, indicating interference by enhanced S7-OP7 vRNA replication (43).

**Figure 6:**
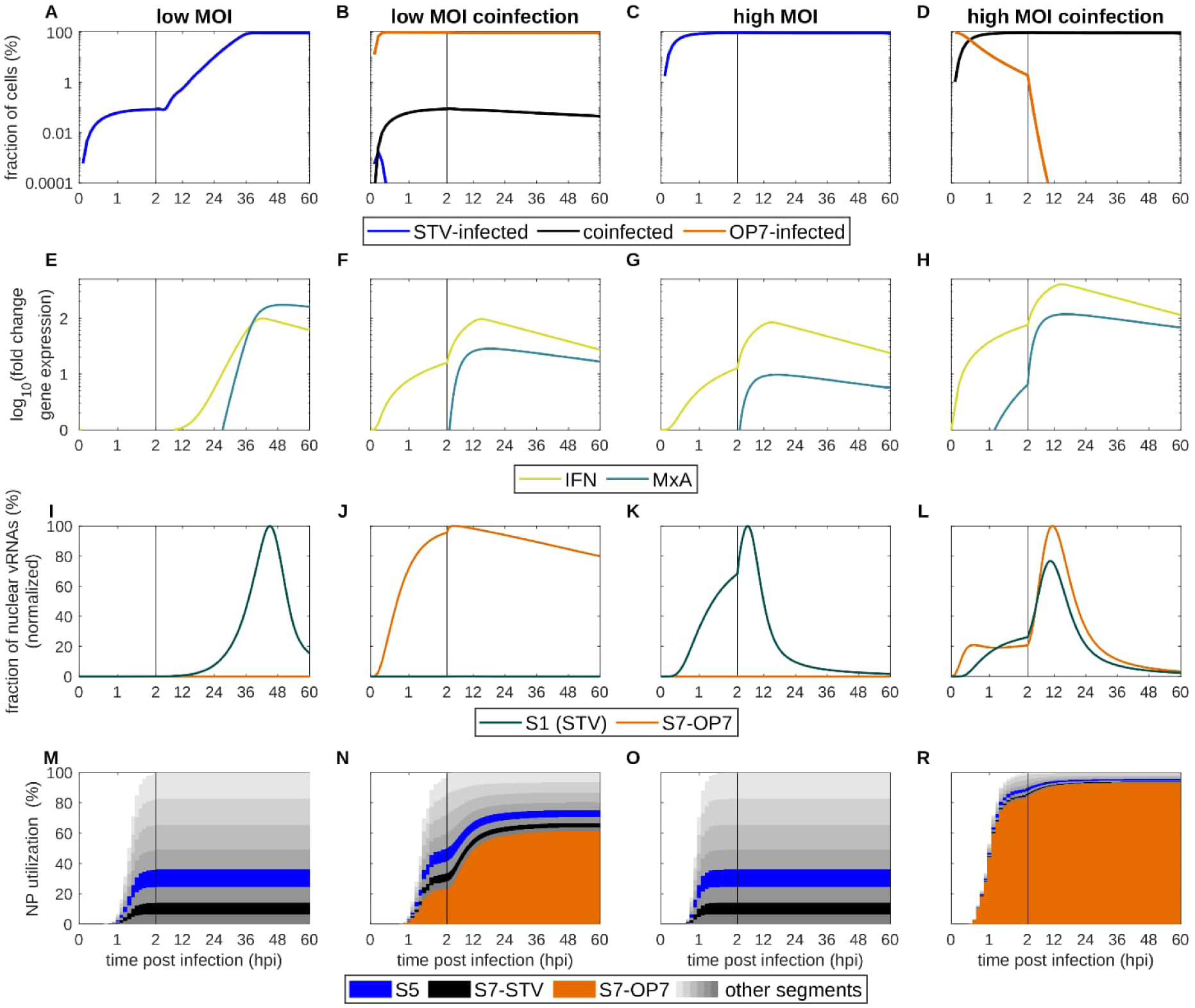
Temporal dynamics of IFN induction and nuclear import of genomic vRNA explain MOI-dependent IFN-mediated antiviral activity. Simulations of STV-only and coinfections for (A) STV MOI of 10^-3^, (B) combined with a low OP7 dose and (C) STV MOI of 3, (D) combined with a high OP7 dose. (A-D) The fraction of STV-infected, coinfected, and OP7-infected cells, (E-H) the activation of the IFN system indicated by the gene expression of IFN-β and MxA, (I-J) the fraction of nuclear vRNAs illustrating the import of vRNAs to and subsequent export from the nucleus, and (M-R) the utilization of viral NP per genome segment, indicating replication interference by OP7, are presented. S1 is used to quantify the import of STV vRNAs as this segment is contained exclusively in STV particles. The vertical line indicates a shift in the scaling on the x-axis at 2 hpi.

In low MOI conditions, a STV-only infection spreads gradually over time, with the majority of cells getting infected between 24 to 36 hpi and all STV vRNAs utilizing similar amounts of viral NP (**Figure 6A+M**). In that time, the IFN response slowly ramps up, but does not appear to manage to prevent the import of STV vRNAs to the nucleus (**Figure 6E+I**). During low MOI coinfection, nearly all cells are infected rapidly by OP7 (**Figure 6B**). Accordingly, the IFN response is activated more quickly, while MxA expression is initiated already at 3 hpi (**Figure 6F**). Thus, in the subsequent phase where most infections would occur in a low MOI STV-only infection, the nuclear import of STV vRNAs is prevented by MxA (**Figure 6A+J**). At the same time, S7-OP7 vRNA nuclear import is unaffected as it entered cells before significant expression of MxA. The residual amounts of STV vRNA entering the nucleus encounter S7-OP7 vRNA, which then occupies about 60 % of the viral NP indicating antiviral replication interference (**Figure 6N**). As a result, only approximately 0.1 % of cells are becoming (co)infected by the STV and the infection is abrogated almost entirely (**Figure 6B**), as indicated by no detectable infectious virus titer (**Figure 5A**, red bar). In high MOI (co)infections, all cells are quickly (co)infected until 2 hpi (**Figure 6C-D**). Consequently, for a STV-only infection, the activation of the IFN response occurs faster than during a low STV MOI infection (**Figure 6G**). However, the nuclear import of STV vRNA precedes the expression of MxA (**Figure 6G+K**). For a high MOI coinfection, IFN-β and MxA expression is increased even more and MxA is synthesized earlier (**Figure 6H**). However, this only reduces, but does not prevent, the import of STV and S7-OP7 vRNA into the nucleus (**Figure 6L**). Subsequently, replication interference, indicated by an occupation of 90 % of viral NP by S7-OP7 vRNA (**Figure 6R**), dominates suppression of STV replication and virus particle release (**Figure 5B**).

In summary, at low MOI coinfection, model simulations suggest that early OP7-induced IFN signaling restricts nuclear import of STV vRNA, while subsequent replication interference reinforces complete suppression of virus replication. At high MOI coinfection, STV vRNA import likely precedes establishment of an effective IFN response, leaving replication interference as the dominant antiviral mechanism of OP7.

## Discussion

In this study, we combined data-driven multiscale modeling with experimental validation to dissect the antiviral mechanisms of the IAV DIP OP7. The model accurately predicted the therapeutic treatment window of OP7, reproduced long-term prophylactic protection following mechanistic refinement, and uncovered an MOI-dependent shift in the relative contributions of IFN-mediated inhibition and replication interference. Experimental validation confirmed these predictions, demonstrating the ability of the model to generate biologically meaningful and experimentally testable hypotheses.

In our implementation of the IFN system, we focus on IFN-β and MxA. While a significantly larger set of different IFNs, ISGs and molecules are involved in the IFN system (75), IFN-β and MxA have been identified as the key players for IAV infection (65, 66). Thus, we take MxA as a representation of the general antiviral activity induced by the IFN response. Next, we compared two competing hypotheses on how MxA inhibits STV replication, i.e., via the formation of dimers that bind to newly synthesized NP removing it from the supply for virus replication and via the binding to incoming vRNAs preventing their nuclear import. In our model simulations, we observe that a prevention of nuclear import describes the experimental data significantly better than an inhibition of replication (**S2 Table**). Therefore, our results support a study by Matzinger *et al.* (73) suggesting an inhibition of nuclear import of genomic vRNA by MxA. In contrast, Nigg *et al.* showed that MxA does not inhibit nuclear import but rather leads to a prevention of viral replication by MxA dimers efficiently binding viral NP (71). However, they introduced MxA expression via plasmids into cells, and did not stimulate a complete IFN response, unlike Matzinger *et al.* who stimulated cells with IFN-α prior to IAV infection, inducing a complete IFN response (73). In this context, Xiao *et al.* proposed that MxA expression alone is insufficient to restrict nuclear import of vRNAs (74). They theorize that either IFN-induced modifications of MxA or other IFN-induced auxiliary molecules are required to facilitate the full antiviral activity of MxA. This line of thinking was also promoted by Haller *et al.*, who were heavily involved in basic MxA research, suggesting the presence of both mechanisms at the same time (66). Our model simulations suggest that in scenarios where the full IFN response is initiated, the prevention of nuclear import is the dominant mechanism of MxA to suppress IAV infection.

Our model predicts that the IFN response alone is capable to reduce infectious virus titers by more than one log in a low MOI coinfection (**Figure 5A**). Yet, in this specific scenario, the larger part of the inhibiting effect is predicted to be induced by replication interference via S7-OP7 vRNA, which reduced titers by an additional two log. However, both effects work in conjunction to reduce STV replication. Thus, while it appears that the majority of the inhibiting effect in the tested IAV infection scenario is exerted by replication interference, the IFN response still plays an important role. This is especially relevant at low STV MOIs, which cause most airborne transmissions in natural settings (39). In addition, we propose that the enhanced IFN response, which is triggered by OP7, mostly impacts low MOI conditions and has no impact in high MOI infections, which could be confirmed experimentally (**Figure 5**). This effect is based on the fact that the initiation of the IFN response, which is a multi-step process involving different signaling cascades, takes a significant amount of time. When STV infection proceeds slowly, as in a low MOI coinfection, the IFN response is established in time to prevent nuclear import of STV vRNAs almost completely (**Figure 6B+F+J**). However, high MOI coinfections progress significantly faster and the IFN response is too slow to prevent nuclear import of STV vRNAs, so that the suppression is almost exclusively mediated by replication interference (**Figure 6D+L+R**).

Another interesting aspect is the interplay between the viral IFN antagonist NS1 and the IFN system. NS1 inhibits the initiation of the IFN response by preventing recognition of incoming vRNAs by PRRs (67, 68). On the other hand, the IFN response inhibits infection and, therefore, NS1 synthesis. Thus, there are two competing processes that fight for predominance during an IAV infection. During STV-only infection at both low and high MOI, NS1 suppresses the IFN response enough for successful virus replication (**Figure 2**). NS1 reaches high levels while IFN-β expression is kept at an intermediate level. A coinfection with OP7 tips this established balance in favor of the IFN response. Providing a high dose of OP7 induces a significantly enhanced expression of IFN-β and MxA. Additionally, it decreases NS1 levels by one log for high MOI conditions (**Figure 2AB-AC**) and even reduces it below the LoD for lower MOIs (**Figure 2Z+AD**). This is caused, firstly, by a reduction of STV vRNA levels in coinfections due to replication interference by S7-OP7 vRNA (**Figure 2H+K+L**). This also affects S8 vRNA encoding for NS1, which leads to a reduction in NS1 synthesis and, in turn, less inhibition of vRNA recognition by PRRs. Secondly, an increased amount of vRNAs is provided by OP7 treatment, which provides more targets for detection by PRRs leading to an enhanced activation of the IFN response. Overall, this showcases the complexity of antiviral mechanisms exerted by OP7. In this context, we recently hypothesized that the enhanced IFN signaling in OP7 coinfected cells is caused by the suppression of the viral IFN antagonist NS1, mediated by replication interference (63), which appears to be mechanistically confirmed through the present modeling work.

When simulating the prophylactic antiviral effect of OP7, a predecessor model initially only predicted an effect until 3 days after OP7 treatment for both low and high STV MOI coinfections (**Figure S3**). However, the experiments by Opitz et al. (63) showed an effect until 7 days, which is supported by previous animal studies indicating the same time windows for conventional DIP treatment (34). We found that this underestimation of the efficacy of prophylactic OP7 treatment is caused by a rapid nuclear export of S7-OP7 vRNAs. As a result, this leads to diminished ability to interfere with STV replication through replication interference. The initial model considered that this export can be mediated even by residual amounts of M1-STV, which are synthesized in OP7-only infections, as the S7-STV segments in the OP7 seed virus are present in low quantities (**Figure 2L**) and produce viral mRNA (**Figure S1**) and proteins. However, a previous study on OP7 has demonstrated that the nuclear export of vRNAs is inhibited in STV/OP7 coinfections (40), which might be caused by M1-OP7 not being able to bind to vRNAs to facilitate nuclear export (43). Implementing this hypothesis into our model enabled us to describe the prophylactic effect of OP7 treatment in low and high MOI conditions (**Figure 3**). The inability of the predecessor model to capture prolonged prophylactic protection highlighted potentially missing biological processes. Incorporating an inhibited nuclear export of vRNAs during OP7-only infection resolved this discrepancy, illustrating how iterative refinement can improve the predictive power of mathematical models.

Overall, our data-driven multiscale model accurately reproduced STV/OP7 co-infection dynamics across diverse experimental conditions and enabled a mechanistic dissection of OP7 antiviral activity. We demonstrate that OP7 suppresses IAV through two complementary mechanisms whose relative contributions are determined by the STV MOI. For low MOI conditions, which likely resemble natural transmission, early IFN-mediated restriction of genomic vRNA nuclear import cooperates with replication interference to achieve complete suppression of virus replication, whereas replication interference dominates at high MOI. Beyond explaining OP7 antiviral activity, the model prospectively predicted therapeutic efficacy and generated mechanistic hypotheses that were subsequently confirmed experimentally, highlighting its value as a predictive systems virology framework. Future extensions incorporating spatial infection dynamics and tissue architecture may further support optimization of dosing strategies and treatment timing while reducing reliance on animal experimentation.

## Methods

The kinetics describing OP7/STV coinfection are based on an intracellular coinfection model published previously (43). We expanded this model by considering the spread of infection on the cell population level based on a study covering a multiscale model of STV and conventional DIP coinfection (57). On the intracellular and cell population level, we included the innate immune response following virus infection represented by the IFN system. In addition to a model expansion, we introduced further changes to the model, i.e., (i) a reduction of the model states to reduce computational cost, (ii) a scaling factor that directly correlates the MOI with the number of incoming STV vRNAs, (iii) and an explicit switch from primary to secondary mRNA transcription.

### Intracellular model

The intracellular model of OP7/STV coinfection is based on a model previously developed in our group (43). This part of the model uses a set of ordinary differential equations describing virus entry, replication and release of virus particles from infected cells. Here, we consider (i) STV particles, (ii) OP7 particles that contain eight S7-OP7 vRNAs, and (iii) particles that contain STV and S7-OP7 vRNAs in a “7+1” configuration. For the complete set of equations and parameters see **S1 Appendix, S3 Table** and **S4 Table**.

To lower the computational burden in our multiscale setup, we reduced the number of model states we consider for simulation of the intracellular level. The original model described all eight STV segments and the S7-OP7 segment individually. As some segments share the exact same dynamics for vRNA replication in the model, i.e., segments 2–6 and 8, we used the vRNA dynamics for segment 5 as representative states that describe all of them at the same time (S1 Appendix, Equations (S13), (S24) and (S29)). This enabled us to reduce the number of model states by ∼30%. We still consider viral mRNA for all segments individually, because the length of the vRNA affects the speed of transcription.

Furthermore, we introduced a scaling factor *F*_MOI_ that correlates the MOI with the actual number of incoming vRNAs on the intracellular level. We calculated this factor by relating the MOI with the concentrations of vRNAs at *t* = 0. In previous studies using this model setup, an offset was applied to align measured vRNA levels at time of infection with the comparatively lower MOIs (43, 56, 57, 59, 60). With this new direct relation, we assume that only ∼2 % of the infecting virus particles are fully infectious, which is in line with the related literature (76). As we define the incoming OP7 via the actual S7-OP7 vRNA concentration measured at *t* = 0, no additional scaling factor is required.

Additionally, we implemented an explicit switch from primary to secondary mRNA transcription, which occurs when vRNAs are replicated to increase the numbers of available templates for transcription. The experimental data describing OP7-only infections enabled us to directly estimate mRNA transcription rates in the absence of vRNA replication (**Figure 2A**, **Figure S1A**). However, we determined significantly lower transcription rates than for STV-only infection and coinfections. Thus, we assumed that there are additional mechanisms that increase mRNA transcription beyond the higher availability of templates. So now, mRNA transcription for segment j is defined as follows:

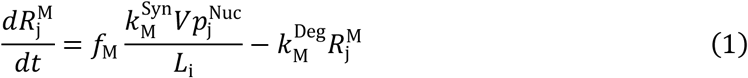

With

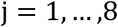

and

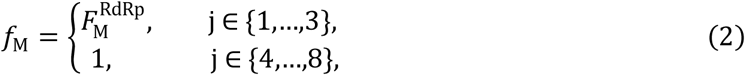

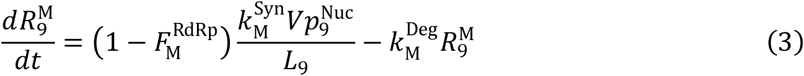

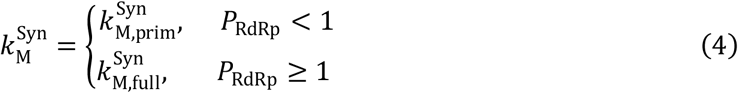

where *f*_M_ is a factor reducing for viral mRNA synthesis RNA-dependent RNA polymerase (RdRp)-related segments and *L*_j_ denotes the length of the viral mRNA related to segment j. S7-OP7 is referred to as segment j = 9. Viral ribonucleoprotein *Vp*^Nuc^, which is vRNA stabilized by RdRp and NP, is used as a template for transcription. The rates 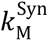 and 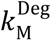 represent the synthesis and degradation of viral mRNA. The concentration of free viral polymerase *P*_RdRp_ is used as an indicator if mRNA transcription is occurring at the speed of primary transcription 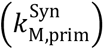 or at full speed 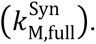.

Lastly, we adjusted the handling of the nuclear export of vRNAs in OP7-only infected cells to accommodate for prophylactic scenarios in which OP7 is administered before STV infection. We assume that, in the absence of vRNA replication, nuclear export is inhibited due to the low amount of available M1-STV leading to a reduction in vRNA binding activity. Therefore, we introduced the binding reduction factor 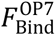 and changed the equation for vRNAs bound by M1 in the nucleus 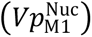 to

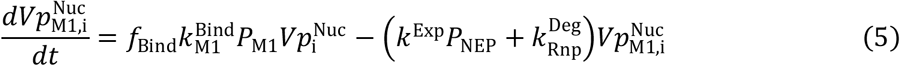

with

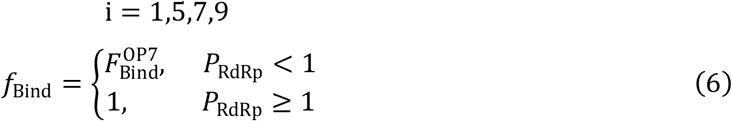

where 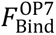 describes the reduced M1-STV binding activity in OP7-only infections. 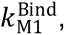 *k*^Exp^ and 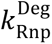 describe the kinetic rates of M1 binding to vRNAs, the export of vRNAs from the nucleus, and the degradation of stabilized vRNAs, respectively. *P*_M1_ and *P*_NEP_ denote the concentrations of M1-STV and the nuclear export protein (NEP, also referred to as NS2), respectively.

### Model of the population level

The cell population level of our coinfection model describes the spread of virus infection between cells. Briefly, a set of ordinary differential equations is combined with integro-partial differential equations to cover virus infection, virus release and the IFN response on the cell population level. The model further describes growth, infection, apoptosis and lysis of uninfected *T*, STV-infected *I*_STV_, OP7-infected *I*_OP7_, and coinfected cells *I*_CO_ as well as the release of STV particles, OP7 particles and IFN-β. For the complete set of equations and parameters see **S1 Appendix**, **S3 Table** and **S4 Table**.

On the population level, we implemented the IFN response using a direct approach. We assume IFN-β is constantly secreted from uninfected cells, which release a baseline amount that keeps the fold change at 1. Infected cells release an increased level of IFN-β to induce protection in other cells. Cells infected with a high dose of OP7 show significantly higher levels of IFN-β expression than STV-only infected cells or when a low OP7 dose is applied (**Figure 2**). Thus, we implemented the release of IFN-β on the cell population as follows:

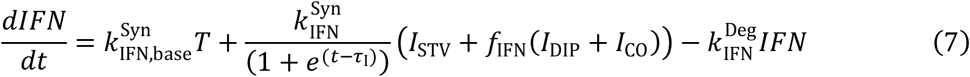

with

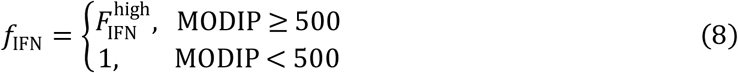

where 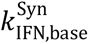 and 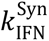 denote the rate of IFN-β release from uninfected and infected cells respectively. 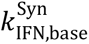 is calibrated to keep a constant baseline level of IFN-β in the system. 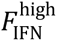 represents a factor that defines the increased release of IFN-β from cells infected with a high OP7 dose. *τ*_I_ denotes the time after which IFN-β production is downregulated by feedback inhibition following infection (77–79). Additionally, IFN-β degradation is described by the parameter 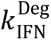 (80).

In contrast to the intracellular level, we only consider (i) STV particles and (ii) OP7 particles that contain eight S7-OP7 vRNAs on the cell population level. Thus, we disregard the release of virus particles that contain STV and S7-OP7 vRNAs in a “7+1” configuration.

### Experimental data and infection conditions

All experimental data used in this study were taken from Opitz *et al*. or were determined according to their protocols (63).

For model simulation, we defined the MOI based on 50% Tissue Culture Infectious Dose (TCID_50_) measurements and for the multiplicity of OP7 (MODIP) we used values based on the intracellular S7-OP7 vRNA concentration measured at *t* = 0 upon infection. Thus, STV MOIs are defined as 0, 10^-3^ and 30. MODIPs were calculated considering eight S7-OP7 vRNAs per OP7 particle and a loss of ∼50 % during the fusion of enveloped viruses (81). The results were rounded to obtain MODIPs of 0, 5 (low dose), 500 (high dose).

For the infection experiment using Ruxo, a different OP7 seed virus was used (63). This led to higher concentrations of S7-OP7 vRNA during infection. We adjusted the initialization for our model simulations shown in **Figure 5 and 6** according to the measured intracellular S7-OP7 vRNA concentration and used MODIPs of 20 (low MOI coinfection) and 2000 (high MOI coinfection).

### Simulation approach and parameter estimation

Model simulations were performed based on a previously published model of STV and DIP coinfection (57). The CVODE routine from SUNDIALS (82) was used to solve model equations numerically on a Linux-based system. Model files and experimental data were handled with the IQM toolbox, formerly known as Systems Biology Toolbox 2 (83), for MatLab (version 9.12.0.1884302, R2022a).

STV-infected and coinfected cells are simulated with the current amount of STV and OP7 particles present on the cell population level at time of infection. To describe cells that are only infected by OP7 and show no replication activity, we simulate the intracellular model once with the initial numbers of OP7 provided and assume all OP7-infected cells behave similarly.

Baseline model parameters were calibrated by fitting the intracellular part of the model to experimental data from an OP7/STV coinfection using infection conditions with high MOIs, i.e., MOI 3 and 30. These single-cycle infections guarantee that the occurrence of multiple waves of infection do not blur the dynamics of the infection cycle of a cell in the experimental data. The NS1 protein level data shown in **Figure 2Y-AD** were not used for parameter estimation as they originate from an altered experimental setup (63). Subsequently, parameters representing the dynamics of IFN-β on the cell population level and the impact of MxA on nuclear import were fitted with the full multiscale model. This includes the parameters *K*_MxA_, 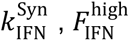 and *x*.

We utilized the global optimization algorithm *GlobalSearch* implemented in MatLab to determine an optimal set of parameters. To determine model fitness, the sum of squared residuals was divided by the standard deviation of the experimental data. All parameter values used for model simulation are shown in **S3 Table** and **S4 Table**.

Using Bayesian inference via an affine invariant ensemble Markov Chain Monte Carlo (MCMC) sampler (84, 85), we obtained the credible intervals of the estimated model parameters in **S1 Table**. Employing the MCMC implementation for MatLab (gwmcmc), we determined the parameter distributions shown in **Figures S5 and S6**. We assumed uniformly distributed priors for the parameters estimated during the initial baseline fit based on the parameter sets determined by the global optimization algorithm. For the parameters related to IFN-β dynamics on the cell population level, we used normally distributed priors around the estimated parameter values as the simulation of the multiscale model takes significantly longer than the intracellular levels alone. Posterior distributions were constructed from 1,000,000 and 76,896 samples when fitting the baseline parameters and the IFN-related parameters to experimental data, respectively.

We applied a burn-in phase to reach an equilibrium distribution before sampling by discarding the first 10% of results obtained. Additionally, we performed a thinning step by retaining every tenth sample to reduce the correlation between individual samples for the baseline parameters. Thus, the resulting parameter distributions contain 89,992 and 76,896 samples for the fits of the baseline parameters and the IFN-related parameters to experimental data, respectively.

### Model prediction

For the prediction of the therapeutic and prophylactic effect of OP7 in **Figure 3**, we simulated infections with a delayed STV infection or OP7 treatment. To achieve a representation of the protective effect determined experimentally in **Figure 3C-D**, we included an inhibited export of vRNAs from the nucleus when cells are only infected by OP7.

To predict the impact of the IFN response on infections at low and high MOIs in **Figure 5**, we simulated the multiscale model for a STV-only infection, a STV/OP7 coinfection and STV/OP7 coinfections in which different sources of the antiviral effect of OP7 were removed. In the setups representing a non-replicating S7-OP7 vRNA, OP7 particles are still infecting the cells, however, incoming S7-OP7 vRNA does not replicate. Thus, they still initiate an IFN response. As specified under **“Infection conditions”**, we adjusted the input of OP7 to match the different concentration of S7-OP7 vRNA in the OP7 seed virus used for this specific experiment.

### Limits of detection

The LoD for NS1 in **Figure 2** was assumed to be similar to the LoD determined for NP and HA protein in Küchler *et al*. (86, 87). The LoD for total particle concentrations in **Figures 2**, **4, S3 and S4** is based on a value of zero determined in the HA assay, calculated to total virus particles/mL according to (88). The LoD for infectious particle concentration in **Figure 4** is based on the lowest possible positive result encountered during the workflow of the TCID_50_ assay (89).

## Supporting information

FullSupplement

## Acknowledgements

We thank Nancy Wynserski for excellent technical assistance.

## Author contributions

Conceptualization, D.R., U.R., S.Y.K.; Data Curation, P.O., J.K.; Formal Analysis, D.R.; Funding Acquisition, U.R., S.Y.K.; Investigation, D.R., P.O., J.K., S.Y.K.; Methodology, D.R., S.Y.K.; Project Administration, D.R., S.Y.K.; Software, D.R.; Supervision, U.R, S.Y.K.; Validation, D.R., P.O., J.K., S.Y.K.; Visualization, D.R.; Writing – Original Draft, D.R.; Writing – Review & Editing, D.R., P.O., U.R., S.Y.K.

