## Supplementary material for "Data-driven multiscale modeling deciphers MOI-dependent dual antiviral mechanisms of OP7": FullSupplement

#### Figures

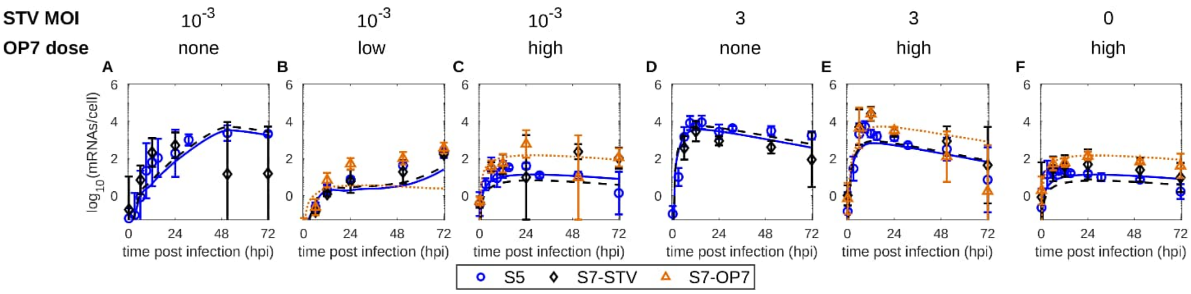

**Figure S1: The coinfection model captures dynamics of viral mRNA synthesis.** Simulations of the STV/OP7 coinfection model fitted to experimental data. (A-F) Cell-specific viral mRNA levels for Calu-3 cells infected with different combinations of STV MOI (influenza A/PR/8/34, H1N1) and OP7 doses. Error bars show the standard deviation of three independent experiments.

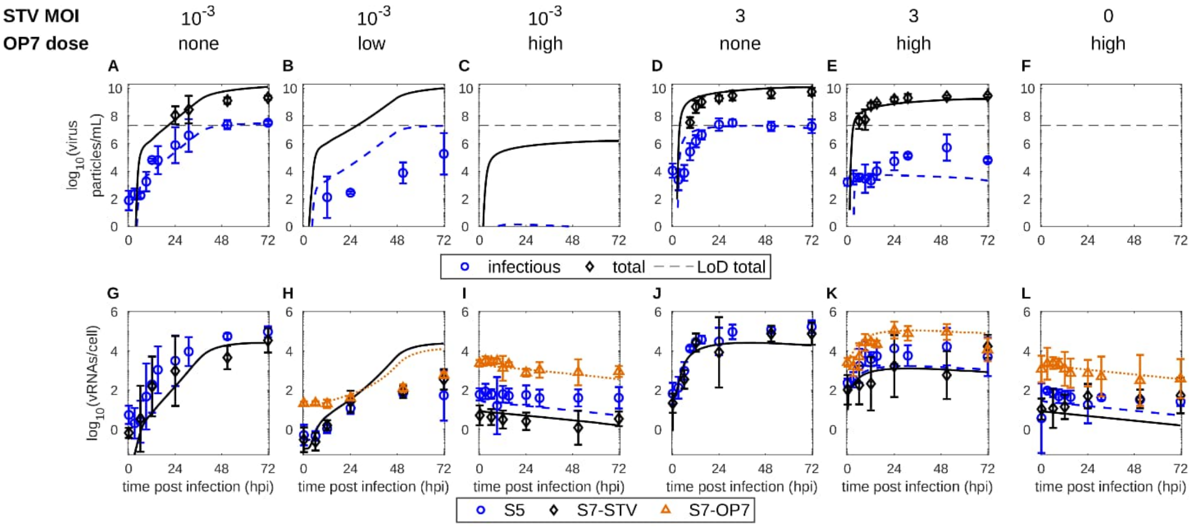

**Figure S2: Simulation results of the coinfection model without an IFN system.** Simulations of the coinfection model without an IFN system fitted to the experimental data. (A-F) Extracellular virus titers and (G-L) cell-specific vRNAs for Calu-3 cells infected with combinations of different STV MOIs (influenza A/PR/8/34, H1N1) and OP7 doses. The black dashed lines indicate the LoD for total virus particles/mL. Error bars show the standard deviation of three independent experiments.

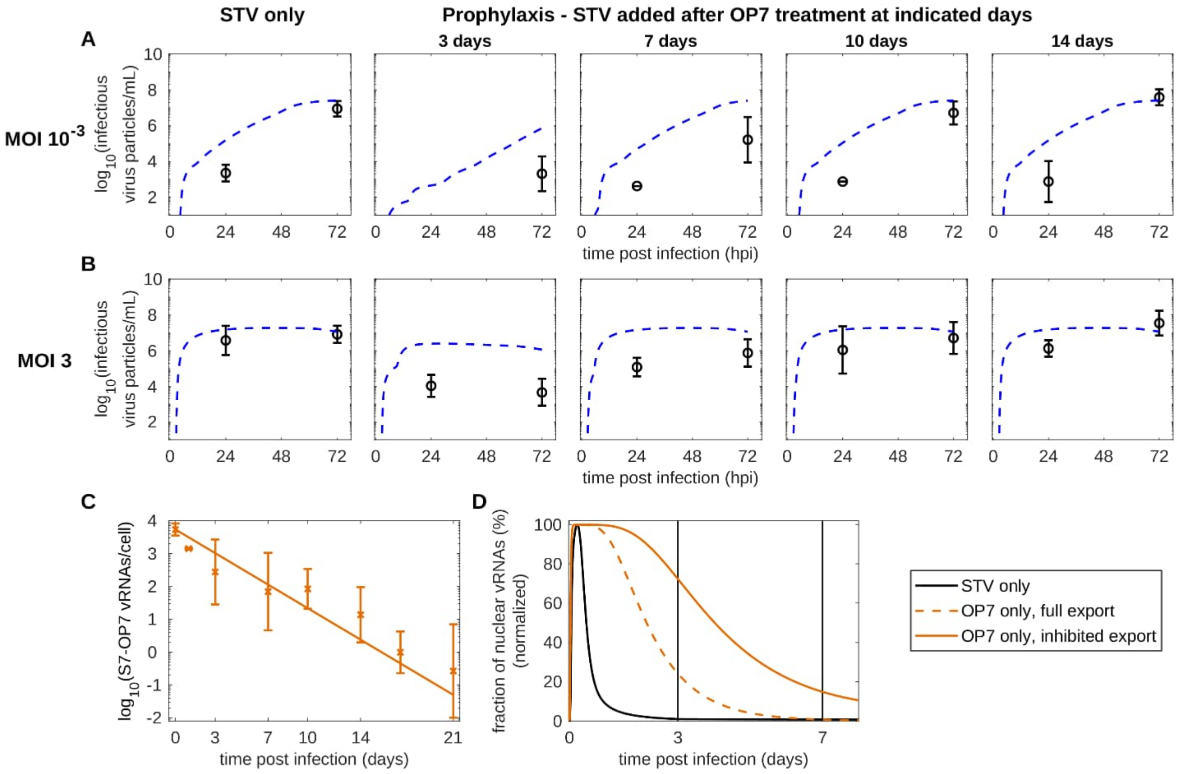

**Figure S3: Initial model prediction of the prophylactic antiviral effect of OP7 and examination of the nuclear export of vRNAs in OP7-only infections.** (A-B) Model prediction and experimental outcome of the prophylactic effect of OP7 added before STV infection using MOIs of $10^{-3}$ and 3. Here, a coinfection model with full export (D) of S7-OP7 vRNA during OP7-only infection was used. (C) Simulation and experimental validation of the degradation of intracellular S7-OP7 vRNA following OP7-only infection. Error bars represent the standard deviation of three independent experiments. (D) Simulation of the export of vRNAs from the nucleus in a STV-only and OP7-only infection using different models. The dashed line represents a full export of S7-OP7 vRNA and the solid line represents an inhibited export of S7-OP7 vRNA due to a low availability of M1-STV.

To uncover the source of the deviation between model simulations and experimental data in (A-B), we first checked if the model predicted the intracellular S7-OP7 vRNA levels over 14 days correctly. If our model would have overestimated the degradation of S7-OP7 vRNA, a prediction of higher virus titers could be explained. However, subsequent experiments confirmed our predicted long-term S7-OP7 vRNA levels (C). Next, we examined if S7-OP7 vRNAs stay in the nucleus after infection, which is where OP7 replication interference with STV replication occurs. We observed that in the initial model simulation, 75% of vRNAs were already exported from the nucleus at 3 days post infection (dpi) and no vRNAs were left at 7 dpi (D). Apparently, this severely weakens the prophylactic effect of OP7 in our simulations

(A-B).

The export of S7-OP7 vRNA in the simulations is mediated by M1-STV derived from the S7-STV present in the OP7 seed virus used for the experimental studies (**Figure 2L**). Although no replication of vRNA occurs in a prophylactic setting, primary transcription, which is the production of viral mRNAs from the incoming vRNAs that infected the cell (1), still takes place (**Figure S1**). This enables the synthesis of viral proteins like M1-STV at very low levels initiating the nuclear export of S7-OP7 vRNAs in the "full export" model (D).

To describe the experimentally observed prophylactic effect, we assumed that such low M1-STV levels, combined with the presence of M1-OP7, would lead to an inhibited nuclear export of vRNAs. M1-OP7 was previously suggested to be defective in binding to vRNAs in the nucleus (2) and might further disturb the export process. A disruption of the nuclear expert of vRNAs was also suggested previously for OP7 (3).

Implementing the hypothesis that the nuclear export of vRNAs is inhibited in OP7-only infected cells (D) enabled us to closely describe the prophylactic effect of OP7 treatment (**Figure 3C+D**). This model extension had no impact on the model simulations in **Figure 2**.

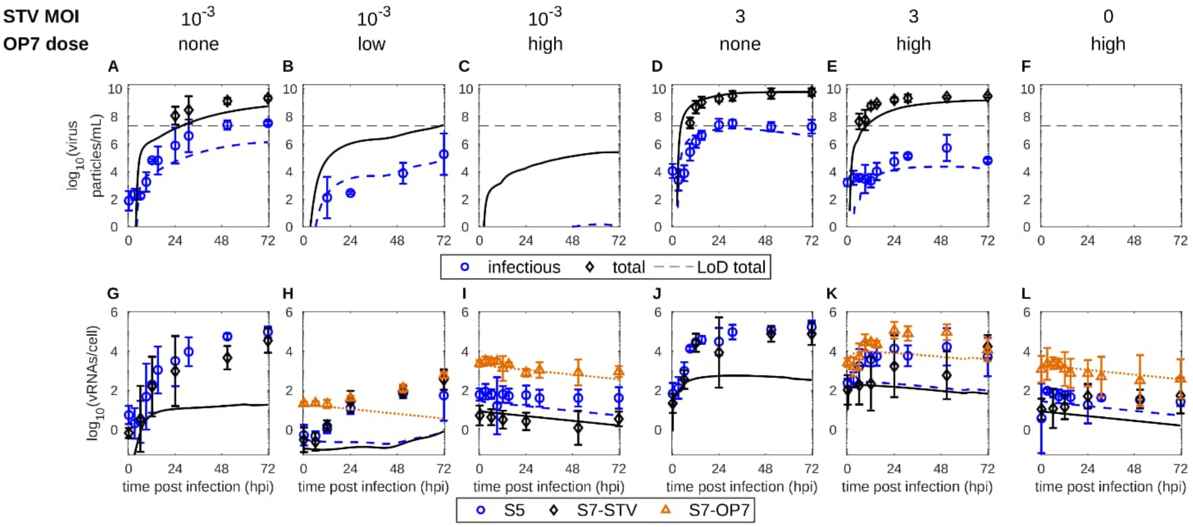

**Figure S4: Simulation results assuming an inhibition of virus replication by MxA.** Simulations of a STV/OP7 coinfection model using the hypothesis that MxA dimers bind to progeny NP to inhibit virus replication. The model was fitted to (A-F) extracellular virus titers and (G-L) cell-specific vRNAs for Calu-3 cells infected using combinations of different STV MOIs (influenza A/PR/8/34, H1N1) and OP7 doses. The black dashed lines indicate the LoD for total virus particles/mL. Error bars show the standard deviation of three independent experiments.

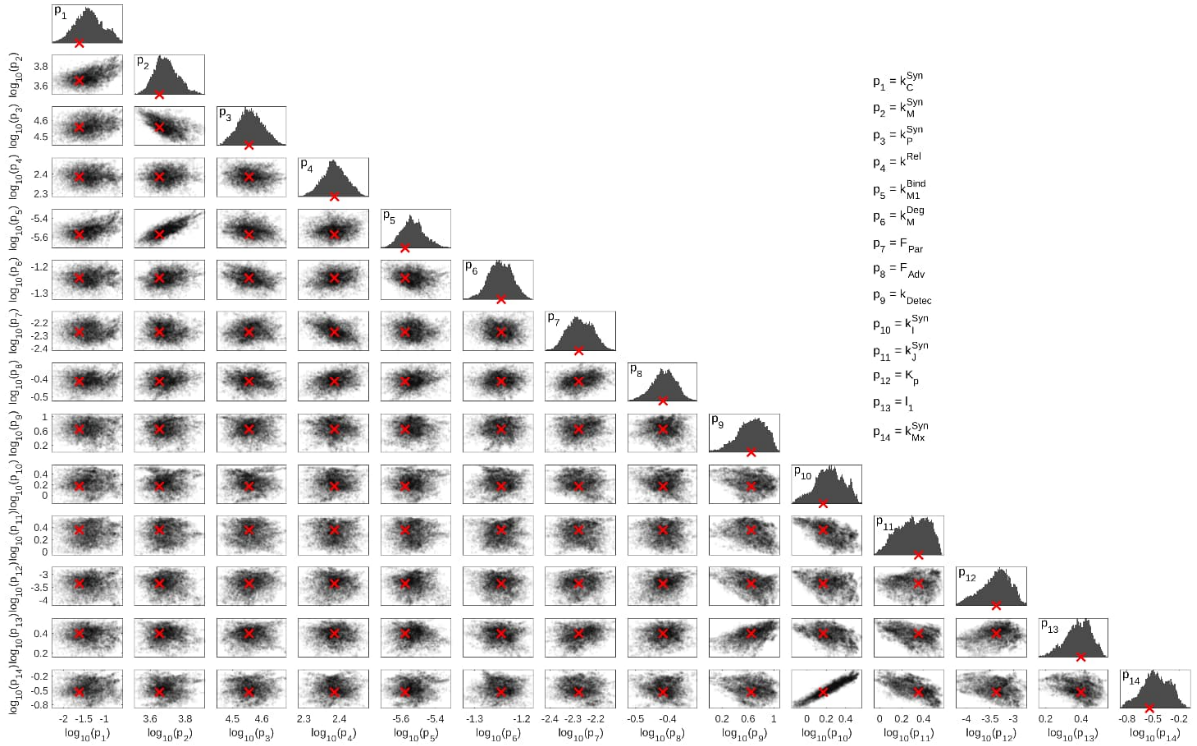

**Figure S5. Parameter distributions and correlations for the calibration to high MOI infection data**. The Histograms show the distributions of each individual estimated parameter. The scatter plots represent the dependency of two respective parameters. Posterior parameter distributions were obtained using an MCMC approach (Bayesian inference). The red X-symbol depicts the parameter value estimated using a global optimization algorithm.

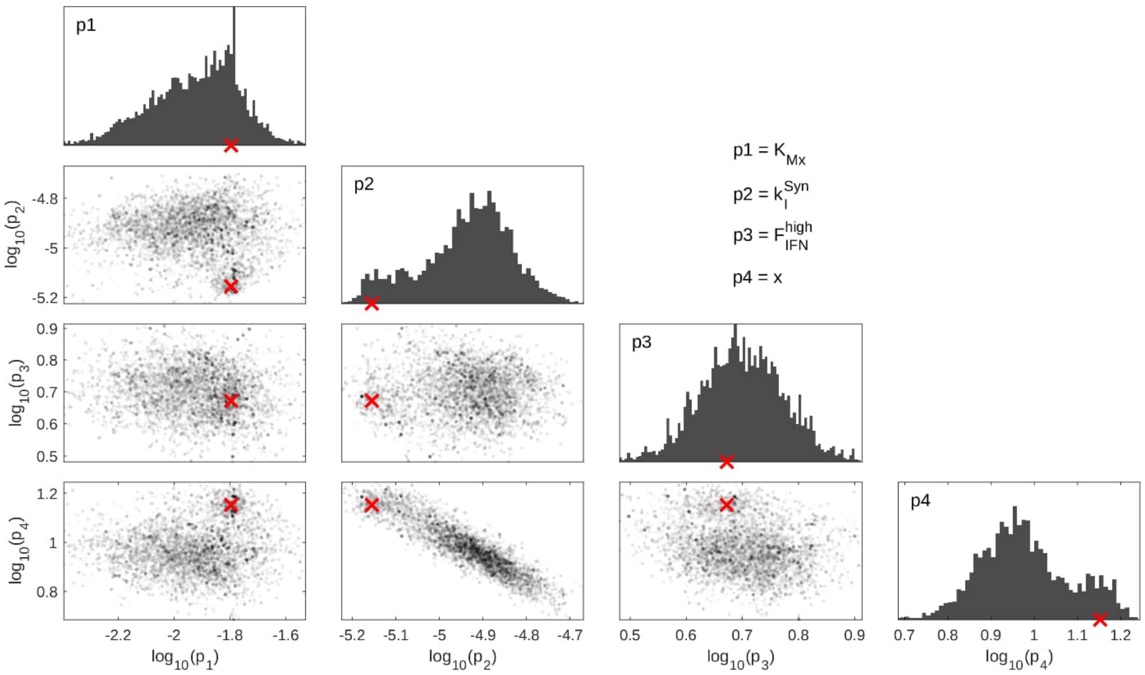

**Figure S6. Parameter distributions and correlations for the calibration of IFN-related parameters.** The histograms show the distributions of each individual parameter. The scatter plots represent the dependency of the two respective parameters. Posterior parameter distributions were determined using an MCMC approach (Bayesian inference). The red X-symbol depicts the parameter value estimated using a global optimization algorithm.

### Tables

**Table S1. Parameters estimated from the experimental data in Figure 1.**

| Parameter | | Value | Credible interval (95%)[a] |
|---|---|---|---|
| $k_{C}^{Syn}$ | ($h^{-1}$) | $2.45\times10^{-2}$ | $(0.09 – 2.02)\times10^{-1}$ |
| $k_{M,full}^{Syn}$ | (nucleotides · $h^{-1}$) | $4.49\times10^{3}$ | $(3.69 – 7.03)\times10^{3}$ |
| $k_{P}^{Syn}$ | (nucleotides · $h^{-1}$) | $3.60\times10^{4}$ | $(3.02 – 4.44)\times10^{4}$ |
| $k^{Rel}$ | (virions · $h^{-1}$) | $2.43\times10^{2}$ | $(2.07 – 2.83)\times10^{2}$ |
| $k_{M1}^{Bind}$ | ($molecules^{-1}$ · $h^{-1}$) | $2.66\times10^{-6}$ | $(2.20 – 4.18)\times10^{-6}$ |
| $k_{M}^{Deg}$ | ($h^{-1}$) | $5.71\times10^{-2}$ | $(5.04 – 6.35)\times10^{-2}$ |
| $F_{Par}$ | (–) | $5.23\times10^{-3}$ | $(4.25 – 6.89)\times10^{-3}$ |
| $F_{Adv}$ | (–) | $3.85\times10^{-1}$ | $(3.24 – 4.54)\times10^{-1}$ |
| $k_{Detec}$ | (molecules · $virions^{-1}$ · $h^{-1}$) | $4.45\times10^{0}$ | $(1.47 – 9.95)\times10^{0}$ |
| $k_{I,in}^{Syn}$ | ($h^{-1}$) | $1.47\times10^{0}$ | $(0.84 – 3.14)\times10^{0}$ |
| $k_{J}^{Syn}$ | ($h^{-1}$) | $2.26\times10^{0}$ | $(1.07 – 3.39)\times10^{0}$ |
| $K_{P}$ | (molecules) | $4.34\times10^{-4}$ | $(0.09 – 1.38)\times10^{-3}$ |
| $\tau_{J}$ | (h) | $2.50\times10^{0}$ | $(1.68 – 3.20)\times10^{0}$ |
| $k_{MxA}^{Syn}$ | ($h^{-1}$) | $2.89\times10^{-1}$ | $(1.62 – 6.75)\times10^{-1}$ |
| $K_{MxA}$ | (molecules) | $1.60\times10^{-2}$ | $(0.54 – 2.23)\times10^{-2}$ |
| $k_{I,ex}^{Syn}$ | ($h^{-1}$) | $7.00\times10^{-6}$ | $(0.68 – 1.77)\times10^{-5}$ |
| $F_{IFN}^{high}$ | (–) | $4.70\times10^{0}$ | $(3.55 – 7.06)\times10^{0}$ |
| $\tau_{I}$ | (h) | $1.42\times10^{1}$ | $(0.61 – 1.54)\times10^{1}$ |

[a] 95% credible intervals were determined from posterior parameter distributions calculated via Bayesian inference (Figures S5 and S6) using a Markov chain Monte Carlo sampler (4).

**Table S2. Table containing the estimated parameter values describing the effect of MxA and the AICc values for the different model hypotheses.**

| **Model version / hypothesis** | $K_{\text{MxA}}$ | $K_{\text{MxA,replication}}$ | **AICc** |
|---|---|---|---|
| Initial model without IFN system | - | - | 1277 |
| Model with IFN hypothesis 1 + 2 | 3000 | 0.0157 | 1141 |
| Model with IFN hypothesis 1 | 0.004 | - | 1820 |
| Model with IFN hypothesis 2 | - | 0.0160 | **1131** |

AICc … corrected Akaike information criterion

**Table S3. Parameters of the intracellular OP7/STV coinfection model.**

| Parameter | Description | Value | Unit | Source |
|---|---|---|---|---|
| $B_{Hi}^{Tot}$ | number of high-affinity binding sites | 150 | sites | (5) |
| $B_{Lo}^{Tot}$ | number of low-affinity binding sites | 1000 | sites | (5) |
| $D_{Rib}$ | distance between two adjacent ribosomes | 160 | nucleotides | (6) |
| $F_{Adv}$ | S7-OP7 cRNA and vRNA replication advantage factor | 0.385 | – | model fit in Fig 2 (see Table S1) |
| $F_{Fus}$ | fraction of fusion-competent virions | 0.51 | – | (7) |
| $F_{MOI}$ | scaling factor correlating MOI with number of incoming vRNAs | 50 | – | calculated based on S1 Data |
| $F_{Par}$ | fraction of infectious STV particles | 0.20 | – | model fit in Fig 2 (see Table S1) |
| $F_{Spl7}$ | fraction of M2-encoding mRNAs | $2.00\times10^{-2}$ | – | (8) |
| $F_{Spl8}$ | fraction of NEP-encoding mRNAs | 0.125 | – | (9) |
| $F_{Bind}^{OP7}$ | reduced M1-STV binding activity in OP7-only infections | $5.50\times10^{-2}$ | – | model fit to data in Figure 3 |
| $F_{M}^{OP7}$ | reduction factor of S7-OP7 mRNA synthesis | 0.9 | – | (2) |
| $F_{M}^{RdRp}$ | reduction factor for RdRp-related viral mRNA synthesis | 0.12 | – | (10) |
| $K_{V^{Rel}}$ | influence of available viral components on virion release | 1 | virions | set to 1 for fast release |
| $k_{Hi}^{Att}$ | attachment to high-affinity binding sites | $8.09\times10^{-2}$ | $site^{-1}\cdot h^{-1}$ | (8) |
| $k_{Lo}^{Att}$ | attachment to low-affinity binding sites | $4.55\times10^{-4}$ | $site^{-1}\cdot h^{-1}$ | (8) |
| $k_{M1}^{Bind}$ | binding of M1-STV to nuclear vRNPs | $2.66\times10^{-6}$ | $molecules^{-1}\cdot h^{-1}$ | model fit in Fig 1 (see Table S1) |
| $k_{NP}^{Bind}$ | binding of NP to RdRp-RNA complexes | 1 | $molecules^{-1}\cdot h^{-1}$ | set to 1 for fast binding of NP |
| $k^{Cplx}$ | formation rate of vRNP complexes | 1 | $molecules^{-7}\cdot h^{-1}$ | (11) |
| $k_{I}^{Deg}$ | degradation of IFN-β | $2.89\times10^{-2}$ | $h^{-1}$ | (12) |
| $k_{J}^{Deg}$ | degradation of JAK | $2.17\times10^{-1}$ | $h^{-1}$ | (13) |
| $k_{M}^{Deg}$ | degradation of mRNA | $5.71\times10^{-2}$ | $h^{-1}$ | model fit in Fig 1 (see Table S1) |

| Parameter | Description | Value | Unit | Source |
|---|---|---|---|---|
| $k_{\mathrm{MxA}}^{\mathrm{Deg}}$ | degradation of MxA | $1.44\times10^{-2}$ | $h^{-1}$ | (14) |
| $k_{\mathrm{Rnp}}^{\mathrm{Deg}}$ | degradation of RNPs | $2.30\times10^{-2}$ | $h^{-1}$ | calculated based on S1 Data |
| $k_{\mathrm{RRdRp}}^{\mathrm{Deg}}$ | degradation of RdRp-RNA complexes | 4.25 | $h^{-1}$ | (7) |
| $k^{\mathrm{En}}$ | endocytosis | 4.8 | $h^{-1}$ | (7) |
| $k_{\mathrm{Hi}}^{\mathrm{Eq}}$ | equilibrium constant of high-affinity sites | $1.13\times10^{-2}$ | $\mathrm{sites}^{-1}$ | (5) |
| $k_{\mathrm{Lo}}^{\mathrm{Eq}}$ | equilibrium constant of low-affinity sites | $8.33\times10^{-5}$ | $\mathrm{sites}^{-1}$ | (5) |
| $k^{\mathrm{Fus}}$ | fusion with endosomes | 3.21 | $h^{-1}$ | (15) |
| $k^{\mathrm{Imp}}$ | nuclear import | 6 | $h^{-1}$ | (16) |
| $K_{\mathrm{MxA}}$ | inhibition of nuclear import by MxA | $1.60\times10^{-2}$ | molecules | model fit in Fig 1 (see Table S1) |
| $K_{\mathrm{P}}$ | inhibition of viral genome recognition by NS1 | $4.34\times10^{-4}$ | molecules | model fit in Fig 1 (see Table S1) |
| $k^{\mathrm{RdRp}}$ | formation of RdRp-complexes | 1 | $\mathrm{molecules}^{-2}\cdot h^{-1}$ | (8) |
| $k^{\mathrm{Rel}}$ | virion release/budding | $2.43\times10^{2}$ | $\mathrm{virions}\cdot h^{-1}$ | model fit in Fig 1 (see Table S1) |
| $k_{\mathrm{C}}^{\mathrm{Syn}}$ | cRNA synthesis | $2.45\times10^{-2}$ | $h^{-1}$ | model fit in Fig 1 (see Table S1) |
| $k_{\mathrm{I,in}}^{\mathrm{Syn}}$ | intracellular IFN-β synthesis | $1.47\times10^{0}$ | $h^{-1}$ | model fit in Fig 1 (see Table S1) |
| $k_{\mathrm{J}}^{\mathrm{Syn}}$ | JAK synthesis | $2.26\times10^{0}$ | $h^{-1}$ | model fit in Fig 1 (see Table S1) |
| $k_{\mathrm{M,full}}^{\mathrm{Syn}}$ | mRNA synthesis | $4.49\times10^{3}$ | $\mathrm{nucleotides}\cdot h^{-1}$ | model fit in Fig 1 (see Table S1) |
| $k_{\mathrm{M,prim}}^{\mathrm{Syn}}$ | primary mRNA synthesis | $1.54\times10^{2}$ | $\mathrm{nucleotides}\cdot h^{-1}$ | model fit to data in Fig S1 |
| $k_{\mathrm{MxA}}^{\mathrm{Syn}}$ | MxA synthesis | $2.89\times10^{-1}$ | $h^{-1}$ | model fit in Fig 1 (see Table S1) |
| $k_{\mathrm{P}}^{\mathrm{Syn}}$ | protein synthesis | $3.60\times10^{4}$ | $\mathrm{nucleotides}\cdot h^{-1}$ | model fit in Fig 1 (see Table S1) |
| $k_{\mathrm{V}}^{\mathrm{Syn}}$ | vRNA synthesis | 1 | $h^{-1}$ | set to 1 to avoid correlation to $k_{\mathrm{C}}^{\mathrm{Syn}}$ |
| $L_1$ | length of segment 1's mRNA | 2320 | nucleotides | (17) |
| $L_2$ | length of segment 2's mRNA | 2320 | nucleotides | (17) |
| $L_3$ | length of segment 3's mRNA | 2211 | nucleotides | (17) |
| $L_4$ | length of segment 4's mRNA | 1757 | nucleotides | (17) |
| $L_5$ | length of segment 5's mRNA | 1540 | nucleotides | (17) |

| Parameter | Description | Value | Unit | Source |
|---|---|---|---|---|
| $L_6$ | length of segment 6’s mRNA | 1392 | nucleotides | (17) |
| $L_7$ | length of segment 7’s unspliced mRNA | 1005 | nucleotides | (17) |
| $L_8$ | length of segment 8’s unspliced mRNA | 868 | nucleotides | (17) |
| $L_9$ | length of OP7 segment 7’s unspliced mRNA | 1005 | nucleotides | same length as STV segment 7 |
| $L_{V,1}$ | length of segment 1’s vRNA and cRNA | 2341 | nucleotides | (17) |
| $L_{V,2}$ | length of segment 2’s vRNA and cRNA | 2341 | nucleotides | (17) |
| $L_{V,3}$ | length of segment 3’s vRNA and cRNA | 2233 | nucleotides | (17) |
| $L_{V,4}$ | length of segment 4’s vRNA and cRNA | 1778 | nucleotides | (17) |
| $L_{V,5}$ | length of segment 5’s vRNA and cRNA | 1565 | nucleotides | (17) |
| $L_{V,6}$ | length of segment 6’s vRNA and cRNA | 1413 | nucleotides | (17) |
| $L_{V,7}$ | length of segment 7’s vRNA and cRNA | 1027 | nucleotides | (17) |
| $L_{V,8}$ | length of segment 8’s vRNA and cRNA | 890 | nucleotides | (17) |
| $L_{V,9}$ | length of OP7 segment 7 vRNA and cRNA | 1027 | nucleotides | same length as STV segment 7 |
| $N_{P_{HA}}$ | number of HA molecules in a virion | 233 | molecules·virion$^{-1}$ | (18) |
| $N_{P_{NA}}$ | number of NA molecules in a virion | 30 | molecules·virion$^{-1}$ | (18) |
| $N_{P_{M1}}$ | number of M1 molecules in a virion | 2700 | molecules·virion$^{-1}$ | (18) |
| $N_{P_{M2}}$ | number of M2 molecules in a virion | 3 | molecules·virion$^{-1}$ | (18) |
| $N_{STV}^{In}$ | average STV segment vRNA in infecting OP7 particles | 0.103 | – | calculated based on S1 Data |
| $N_{S7\text{-}STV}^{In}$ | average S7-STV segment vRNA in infecting OP7 particles | 3.20×10$^{-2}$ | – | calculated based on S1 Data |
| $N_{S7\text{-}OP7}^{In}$ | average S7-OP7 segment vRNA in infecting OP7 particles | 7.293 | – | calculated based on S1 Data |
| $N_{M1}^{Nuc}$ | nucleotides bound by one M1 molecule | 200 | nucleotides | (19) |
| $N_{NP}^{Nuc}$ | nucleotides bound by one NP molecule | 24 | nucleotides | (20) |
| $N_{OP7}^{Par}$ | number of S7-OP7 vRNPs in OP7 particle | 8 | molecules | analogous to STV particles |

| Parameter | Description | Value | Unit | Source |
|---|---|---|---|---|
| $\tau_J$ | time after which JAK production is downregulated by feedback inhibition | 2.5 | h | model fit in Fig 1 (see Table S1) |

**Table S4. Parameters of the cell population model.**

| Parameter | Description | Value | Unit | Source |
|---|---|---|---|---|
| $\tau_{\mathrm{SIE}}$ | time for superinfection exclusion | 3 | h | (15) |
| $\mu_{\mathrm{Max}}$ | maximum cell growth rate | $3.00\times10^{-2}$ | $h^{-1}$ | (21) |
| $B_{\mathrm{Hi}}^{\mathrm{Tot}}$ | number of high-affinity binding sites | 150 | $\mathrm{sites}\cdot\mathrm{cell}^{-1}$ | (5) |
| $B_{\mathrm{Lo}}^{\mathrm{Tot}}$ | number of low-affinity binding sites | 1000 | $\mathrm{sites}\cdot\mathrm{cell}^{-1}$ | (5) |
| $F_{\mathrm{IFN}}^{\mathrm{high}}$ | factor defining the increased release of IFN-β from cells infected with a high OP7 dose | 4.7 | - | model fit in Fig 1 (see Table S1) |
| $F_{\mathrm{Inf}}$ | ratio of infected cells to fused virions | 1 | $\mathrm{cells}\cdot\mathrm{virion}^{-1}$ | (8) |
| $K_{\mathrm{I}}$ | maximum apoptosis rate of infected cell | 0.27 | $h^{-1}$ | (15) |
| $k_{\mathrm{T}}^{\mathrm{Apo}}$ | apoptosis rate of uninfected cells | $1.2\times10^{-2}$ | $h^{-1}$ | (15) |
| $k_{\mathrm{c,Hi}}^{\mathrm{Att}}$ | attachment to high-affinity binding sites | $3.32\times10^{-8}$ | $\mathrm{mL}\cdot\mathrm{sites}^{-1}\cdot h^{-1}$ | (8) |
| $k_{\mathrm{c,Lo}}^{\mathrm{Att}}$ | attachment to low-affinity binding sites | $1.85\times10^{-10}$ | $\mathrm{mL}\cdot\mathrm{sites}^{-1}\cdot h^{-1}$ | (8) |
| $k_{\mathrm{I}}^{\mathrm{Deg}}$ | degradation of IFN-β | $2.89\times10^{-2}$ | $h^{-1}$ | (12) |
| $k_{\mathrm{V}}^{\mathrm{Deg}}$ | degradation/clearance of infectious virions | $5.0\times10^{-2}$ | $h^{-1}$ | adjusted to infectious titer reduction observed in experiments (S1 Data) |
| $k^{\mathrm{En}}$ | endocytosis | 4.8 | $h^{-1}$ | (7) |
| $k_{\mathrm{c,Hi}}^{\mathrm{Eq}}$ | equilibrium constant of high-affinity sites | $4.48\times10^{-9}$ | $\mathrm{mL}\cdot\mathrm{sites}^{-1}$ | (5) |
| $k_{\mathrm{c,Lo}}^{\mathrm{Eq}}$ | equilibrium constant of low-affinity sites | $3.32\times10^{-11}$ | $\mathrm{mL}\cdot\mathrm{sites}^{-1}$ | (5) |
| $k^{\mathrm{Fus}}$ | fusion with endosomes | 3.21 | $h^{-1}$ | (15) |
| $k^{\mathrm{Lys}}$ | lysis of apoptotic cells | 0.16 | $h^{-1}$ | (15) |
| $k_{\mathrm{I,base}}^{\mathrm{Syn}}$ | IFN-β synthesis rate of uninfected cells | $1.40\times10^{-8}$ | $h^{-1}$ | calibrated to maintain baseline level of IFN-β |
| $k_{\mathrm{I,ex}}^{\mathrm{Syn}}$ | IFN-β synthesis rate of infected cells | $7.00\times10^{-6}$ | $h^{-1}$ | model fit in Fig 1 (see Table S1) |
| $T_{\mathrm{Max}}$ | maximum cell concentration | $1\times10^{7}$ | $\mathrm{cells}\cdot\mathrm{mL}^{-1}$ | (15) |
| $\tau_{\mathrm{Apo}}$ | time after cell infection at which the rate of virus-induced apoptosis reaches its half-maximum | 60 | h | adjusted to cell survival times observed in experiments (S1 Data) |

| Parameter | Description | Value | Unit | Source |
|---|---|---|---|---|
| $\tau_{I}$ | time after which IFN-β production is downregulated by feedback inhibition | 14.2 | h | model fit in Fig 1 (see Table S1) |
| $v_{Apo}$ | distribution factor of the virus-induced apoptosis rate | 1.7 | $h^{-1}$ | (15) |

### Appendix

**S1 Appendix. Multiscale model of OP7/STV coinfection.**

**Intracellular level**

$$\frac{dV^{\mathrm{Ex}}}{dt} = k_{\mathrm{Hi}}^{\mathrm{Dis}} V_{\mathrm{Hi}}^{\mathrm{Att}} + k_{\mathrm{Lo}}^{\mathrm{Dis}} V_{\mathrm{Lo}}^{\mathrm{Att}} - \left(k_{\mathrm{Hi}}^{\mathrm{Att}} B_{\mathrm{Hi}} + k_{\mathrm{Lo}}^{\mathrm{Att}} B_{\mathrm{Lo}}\right) V^{\mathrm{Ex}} \tag{1}$$

$$\frac{dD^{\mathrm{Ex}}}{dt} = k_{\mathrm{Hi}}^{\mathrm{Dis}} D_{\mathrm{Hi}}^{\mathrm{Att}} + k_{\mathrm{Lo}}^{\mathrm{Dis}} D_{\mathrm{Lo}}^{\mathrm{Att}} - \left(k_{\mathrm{Hi}}^{\mathrm{Att}} B_{\mathrm{Hi}} + k_{\mathrm{Lo}}^{\mathrm{Att}} B_{\mathrm{Lo}}\right) D^{\mathrm{Ex}} \tag{2}$$

$$\text{with } B_{\mathrm{n}} = B_{\mathrm{n}}^{\mathrm{Tot}} - V_{\mathrm{n}}^{\mathrm{Att}}, \ \mathrm{n} \in \{\mathrm{Hi,Lo}\} \tag{3}$$

$$\text{and } k_{\mathrm{n}}^{\mathrm{Dis}} = \frac{k_{\mathrm{n}}^{\mathrm{Att}}}{k_{\mathrm{n}}^{\mathrm{Eq}}} \tag{4}$$

$$\frac{dV_{\mathrm{n}}^{\mathrm{Att}}}{dt} = k_{\mathrm{n}}^{\mathrm{Att}} B_{\mathrm{n}} V^{\mathrm{Ex}} - \left(k_{\mathrm{n}}^{\mathrm{Dis}} + k^{\mathrm{En}}\right) V_{\mathrm{n}}^{\mathrm{Att}} \tag{5}$$

$$\frac{dD_{\mathrm{n}}^{\mathrm{Att}}}{dt} = k_{\mathrm{n}}^{\mathrm{Att}} B_{\mathrm{n}} D^{\mathrm{Ex}} - \left(k_{\mathrm{n}}^{\mathrm{Dis}} + k^{\mathrm{En}}\right) D_{\mathrm{n}}^{\mathrm{Att}} \tag{6}$$

$$\frac{dV^{\mathrm{En}}}{dt} = k^{\mathrm{En}}\left(V_{\mathrm{Hi}}^{\mathrm{Att}} + V_{\mathrm{Lo}}^{\mathrm{Att}}\right) - \left(k^{\mathrm{Fus}} + k_{\mathrm{En}}^{\mathrm{Deg}}\right) V^{\mathrm{En}} \tag{7}$$

$$\frac{dD^{\mathrm{En}}}{dt} = k^{\mathrm{En}}\left(D_{\mathrm{Hi}}^{\mathrm{Att}} + D_{\mathrm{Lo}}^{\mathrm{Att}}\right) - \left(k^{\mathrm{Fus}} + k_{\mathrm{En}}^{\mathrm{Deg}}\right) D^{\mathrm{En}} \tag{8}$$

$$\text{with } k_{\mathrm{En}}^{\mathrm{Deg}} = \frac{1 - F_{\mathrm{Fus}}}{F_{\mathrm{Fus}}} k^{\mathrm{Fus}}, \ 0 < F_{\mathrm{Fus}} \leq 1 \tag{9}$$

<u>Virus entry:</u> $V^{\mathrm{Ex}}$ and $D^{\mathrm{Ex}}$ represent extracellular STV and OP7 particles, respectively, that bind to free high- and low-affinity binding sites ($B_{\mathrm{n}}$) on the cell membrane (5, 7). We assume that the different particles follow the same entry mechanism. Virions attached to binding sites ($V_{\mathrm{n}}^{\mathrm{Att}}$, $D_{\mathrm{n}}^{\mathrm{Att}}$) either dissociate from the cell membrane or undergo receptor-mediated endocytosis with rates $k_{\mathrm{n}}^{\mathrm{Dis}}$ and $k^{\mathrm{En}}$, respectively. The enveloped virus particles ($V^{\mathrm{En}}$, $D^{\mathrm{En}}$) can transfer their viral genome into the cytoplasm or are degraded in lysosomes.

$$\frac{dV^{\mathrm{Cyt}}}{dt} = k^{\mathrm{Fus}} V^{\mathrm{En}} - k^{\mathrm{Imp}} V^{\mathrm{Cyt}} \tag{10}$$

$$\frac{dD^{\mathrm{Cyt}}}{dt} = k^{\mathrm{Fus}} D^{\mathrm{En}} - k^{\mathrm{Imp}} D^{\mathrm{Cyt}} \tag{11}$$

$$\frac{dVp_1^{\mathrm{Nuc}}}{dt} = \frac{k^{\mathrm{Imp}} V^{\mathrm{Cyt}}}{1 + \frac{G_{\mathrm{MxA}}}{K_{\mathrm{MxA}}}} + k_{\mathrm{NP}}^{\mathrm{Bind}} P_{\mathrm{NP}} R_{\mathrm{RdRp,1}}^{\mathrm{V}} - \left(f_{\mathrm{Bind}} k_{\mathrm{M1}}^{\mathrm{Bind}} P_{\mathrm{M1}} + k_{\mathrm{Rnp}}^{\mathrm{Deg}}\right) Vp_1^{\mathrm{Nuc}} \tag{12}$$

$$\frac{dVp_5^{\mathrm{Nuc}}}{dt} = \frac{k^{\mathrm{Imp}}}{1 + \frac{G_{\mathrm{MxA}}}{K_{\mathrm{MxA}}}}\left(V^{\mathrm{Cyt}} + N_{\mathrm{STV}}^{\mathrm{In}} D^{\mathrm{Cyt}}\right) + k_{\mathrm{NP}}^{\mathrm{Bind}} P_{\mathrm{NP}} R_{\mathrm{RdRp,5}}^{\mathrm{V}} - \left(f_{\mathrm{Bind}} k_{\mathrm{M1}}^{\mathrm{Bind}} P_{\mathrm{M1\text{-}STV}} + k_{\mathrm{Rnp}}^{\mathrm{Deg}}\right) Vp_5^{\mathrm{Nuc}} \tag{13}$$

$$\frac{dVp_7^{\mathrm{Nuc}}}{dt} = \frac{k^{\mathrm{Imp}}}{1 + \frac{G_{\mathrm{MxA}}}{K_{\mathrm{MxA}}}}\left(V^{\mathrm{Cyt}} + N_{\mathrm{S7\text{-}STV}}^{\mathrm{In}} D^{\mathrm{Cyt}}\right) + k_{\mathrm{NP}}^{\mathrm{Bind}} P_{\mathrm{NP}} R_{\mathrm{RdRp,7}}^{\mathrm{V}} - \left(f_{\mathrm{Bind}} k_{\mathrm{M1}}^{\mathrm{Bind}} P_{\mathrm{M1\text{-}STV}} + k_{\mathrm{Rnp}}^{\mathrm{Deg}}\right) Vp_7^{\mathrm{Nuc}} \tag{14}$$

$$\frac{dVp_9^{\mathrm{Nuc}}}{dt} = \frac{k^{\mathrm{Imp}}}{1 + \frac{G_{\mathrm{MxA}}}{K_{\mathrm{MxA}}}}\left(V^{\mathrm{Cyt}} + N_{\mathrm{S7\text{-}OP7}}^{\mathrm{In}} D^{\mathrm{Cyt}}\right) + k_{\mathrm{NP}}^{\mathrm{Bind}} P_{\mathrm{NP}} R_{\mathrm{RdRp,9}}^{\mathrm{V}} - \left(f_{\mathrm{Bind}} k_{\mathrm{M1}}^{\mathrm{Bind}} P_{\mathrm{M1\text{-}STV}} + k_{\mathrm{Rnp}}^{\mathrm{Deg}}\right) Vp_9^{\mathrm{Nuc}} \tag{15}$$

$$f_{\mathrm{Bind}} = \begin{cases} F_{\mathrm{Bind}}^{\mathrm{OP7}}, & P_{\mathrm{RdRp}} < 1 \\ 1, & P_{\mathrm{RdRp}} \geq 1 \end{cases} \tag{16}$$

$$\frac{dR_{\mathrm{RdRp,j}}^{\mathrm{C}}}{dt} = k_{\mathrm{C}}^{\mathrm{Syn}} P_{\mathrm{RdRp}} Vp_{\mathrm{j}}^{\mathrm{Nuc}} - \left(k_{\mathrm{NP}}^{\mathrm{Bind}} P_{\mathrm{NP}} + k_{\mathrm{RRdRp}}^{\mathrm{Deg}}\right) R_{\mathrm{RdRp,j}}^{\mathrm{C}} \tag{17}$$

$$\frac{dR_{\mathrm{RdRp,9}}^{\mathrm{C}}}{dt} = (1 + F_{\mathrm{Adv}}) k_{\mathrm{C}}^{\mathrm{Syn}} P_{\mathrm{RdRp}} Vp_9^{\mathrm{Nuc}} - \left(k_{\mathrm{NP}}^{\mathrm{Bind}} P_{\mathrm{NP}} + k_{\mathrm{RRdRp}}^{\mathrm{Deg}}\right) R_{\mathrm{RdRp,9}}^{\mathrm{C}} \tag{18}$$

$$\frac{dCp_{\mathrm{i}}}{dt} = k_{\mathrm{NP}}^{\mathrm{Bind}} P_{\mathrm{NP}} R_{\mathrm{RdRp,i}}^{\mathrm{C}} - k_{\mathrm{Rnp}}^{\mathrm{Deg}} Cp_{\mathrm{i}} \tag{19}$$

$$\frac{dR_{\mathrm{RdRp,j}}^{\mathrm{V}}}{dt} = k_{\mathrm{V}}^{\mathrm{Syn}} P_{\mathrm{RdRp}} Cp_{\mathrm{j}} - \left(k_{\mathrm{NP}}^{\mathrm{Bind}} P_{\mathrm{NP}} + k_{\mathrm{RRdRp}}^{\mathrm{Deg}}\right) R_{\mathrm{RdRp,j}}^{\mathrm{V}} \tag{20}$$

$$\frac{dR_{\mathrm{RdRp,9}}^{\mathrm{V}}}{dt} = (1 + F_{\mathrm{Adv}}) k_{\mathrm{V}}^{\mathrm{Syn}} P_{\mathrm{RdRp}} Cp_9 - \left(k_{\mathrm{NP}}^{\mathrm{Bind}} P_{\mathrm{NP}} + k_{\mathrm{RRdRp}}^{\mathrm{Deg}}\right) R_{\mathrm{RdRp,9}}^{\mathrm{V}} \tag{21}$$

$$\frac{dVp_{\mathrm{M1,i}}^{\mathrm{Nuc}}}{dt} = f_{\mathrm{Bind}} k_{\mathrm{M1}}^{\mathrm{Bind}} P_{\mathrm{M1}} Vp_{\mathrm{i}}^{\mathrm{Nuc}} - \left(k^{\mathrm{Exp}} P_{\mathrm{NEP}} + k_{\mathrm{Rnp}}^{\mathrm{Deg}}\right) Vp_{\mathrm{M1,i}}^{\mathrm{Nuc}} \tag{22}$$

$$\frac{dVp_{\mathrm{M1,1}}^{\mathrm{Cyt}}}{dt} = k^{\mathrm{Exp}} P_{\mathrm{NEP}} Vp_{\mathrm{M1,1}}^{\mathrm{Nuc}} - k_{\mathrm{Rnp}}^{\mathrm{Deg}} Vp_{\mathrm{M1,1}}^{\mathrm{Cyt}} - k^{\mathrm{Cplx}}\left(Vp_{\mathrm{M1,7}}^{\mathrm{Cyt}} + Vp_{\mathrm{M1,9}}^{\mathrm{Cyt}}\right) Vp_{\mathrm{M1,1}}^{\mathrm{Cyt}} 6 Vp_{\mathrm{M1,5}}^{\mathrm{Cyt}} \tag{23}$$

$$\frac{dVp_{\mathrm{M1,5}}^{\mathrm{Cyt}}}{dt} = k^{\mathrm{Exp}} P_{\mathrm{NEP}} Vp_{\mathrm{M1,5}}^{\mathrm{Nuc}} - k_{\mathrm{Rnp}}^{\mathrm{Deg}} Vp_{\mathrm{M1,5}}^{\mathrm{Cyt}} - k^{\mathrm{Cplx}}\left(Vp_{\mathrm{M1,7}}^{\mathrm{Cyt}} + Vp_{\mathrm{M1,9}}^{\mathrm{Cyt}}\right) Vp_{\mathrm{M1,1}}^{\mathrm{Cyt}} 6 Vp_{\mathrm{M1,5}}^{\mathrm{Cyt}} \tag{24}$$

$$\frac{dVp_{\mathrm{M1,7}}^{\mathrm{Cyt}}}{dt} = k^{\mathrm{Exp}} P_{\mathrm{NEP}} Vp_{\mathrm{M1,7}}^{\mathrm{Nuc}} - k_{\mathrm{Rnp}}^{\mathrm{Deg}} Vp_{\mathrm{M1,7}}^{\mathrm{Cyt}} - k^{\mathrm{Cplx}} Vp_{\mathrm{M1,7}}^{\mathrm{Cyt}} Vp_{\mathrm{M1,1}}^{\mathrm{Cyt}} 6 Vp_{\mathrm{M1,5}}^{\mathrm{Cyt}} \tag{25}$$

$$\frac{dVp_{\mathrm{M1,9}}^{\mathrm{Cyt}}}{dt} = k^{\mathrm{Exp}} P_{\mathrm{NEP}} Vp_{\mathrm{M1,9}}^{\mathrm{Nuc}} - k_{\mathrm{Rnp}}^{\mathrm{Deg}} Vp_{\mathrm{M1,9}}^{\mathrm{Cyt}} - k^{\mathrm{Cplx}} Vp_{\mathrm{M1,9}}^{\mathrm{Cyt}} Vp_{\mathrm{M1,1}}^{\mathrm{Cyt}} 6 Vp_{\mathrm{M1,5}}^{\mathrm{Cyt}}$$
$$-k^{\mathrm{Rel}} \frac{Vp_{\mathrm{M1,9}}^{\mathrm{Cyt}}}{V_{\mathrm{Cplx}}^{\mathrm{Cyt}} + D_{\mathrm{Cplx}}^{\mathrm{Cyt}} + Vp_{\mathrm{M1,9}}^{\mathrm{Cyt}} + K_{\mathrm{V^{Rel}}}} \prod_{\mathrm{p}} \frac{P_{\mathrm{p}}}{P_{\mathrm{p}} + N_{P_{\mathrm{p}}} K_{\mathrm{V^{Rel}}}} \tag{26}$$

with the viral segments i = 1,5,7,9 ; j = 1,5,7 and p ∈ {HA, NA, M1, M2}

<u>Virus replication:</u> Subsequently, viral ribonucleoproteins (vRNPs), which are viral genomic

RNAs (vRNAs) stabilized by RdRp and NP, in the cytoplasm ($V^{\mathrm{Cyt}}$ and $D^{\mathrm{Cyt}}$) are imported into the nucleus with rate $k^{\mathrm{Imp}}$. This import can be inhibited by myxovirus resistance protein 1 (MxA) represented by $G_{\mathrm{MxA}}$. $K_{\mathrm{MxA}}$ describes the strength of this import inhibition. The composition of the OP7 seed virus is considered via the average amount of vRNAs per virion of S7-STV, S7-OP7 and the other STV segments ($N^{\mathrm{In}}_{\mathrm{S7\text{-}STV}}$, $N^{\mathrm{In}}_{\mathrm{S7\text{-}OP7}}$, $N^{\mathrm{In}}_{\mathrm{STV}}$). In the nucleus, the vRNPs of different viral genome segments ($Vp^{\mathrm{Nuc}}_{\mathrm{i}}$) are used as templates for virus replication. We assume that segments 2, 3, 4, 5, 6 and 8 behave similarly and represent them by segment 5. Thus, the simulated segments are i = 1,5,7,9 with i = 9 representing S7-OP7. Complementary RNA (cRNA) $R^{\mathrm{C}}_{\mathrm{RdRp,i}}$ is transcribed from nuclear vRNPs by viral RNA-dependent-RNA-polymerase (RdRp) $P_{\mathrm{RdRp}}$ and subsequently stabilized by binding nucleoproteins (NP) $P_{\mathrm{NP}}$ to form complementary ribonucleoproteins (cRNP) $Cp_{\mathrm{i}}$. Then, progeny vRNA is transcribed from the cRNP templates by RdRp and bound by NP forming $R^{\mathrm{V}}_{\mathrm{RdRp,i}}$ and $Vp^{\mathrm{Nuc}}_{\mathrm{i}}$, respectively. We assume that S7-OP7 cRNA and vRNA show an enhanced replication compared to the STV segments, which is implemented in Equations (18) and (21) via the (advantage) factor $F_{\mathrm{Adv}}$ . The viral matrix protein 1 (M1) derived from S7-STV (M1-STV) $P_{\mathrm{M1\text{-}STV}}$ can bind to vRNPs in the nucleus to form vRNP-M1 complexes ($Vp^{\mathrm{Nuc}}_{\mathrm{M1,i}}$), which are replication-incompetent. We assume that the M1 derived from S7-OP7 (M1-OP7) $P_{\mathrm{M1\text{-}OP7}}$ is defective and does not bind to vRNPs in the nucleus. $F^{\mathrm{OP7}}_{\mathrm{Bind}}$ describes the reduced M1-STV binding activity in OP7-only infections, which is determined by the absence of any synthesis of viral RdRp. For the complex formation of $Vp^{\mathrm{Cyt}}_{\mathrm{M1,i}}$, we multiply $Vp^{\mathrm{Cyt}}_{\mathrm{M1,5}}$ by six to represent the other segments it represents. Then, the nuclear export protein (NEP) $P_{\mathrm{NEP}}$ can bind to the vRNP-M1 complexes, which enables their export to the cytoplasm where they are referred to as $Vp^{\mathrm{Cyt}}_{\mathrm{M1,i}}$.

$$\frac{dR^{\mathrm{M}}_{\mathrm{j}}}{dt} = f_{\mathrm{M}}\,\frac{k^{\mathrm{Syn}}_{\mathrm{M}}Vp^{\mathrm{Nuc}}_{\mathrm{j}}}{L_{\mathrm{i}}} - k^{\mathrm{Deg}}_{\mathrm{M}}R^{\mathrm{M}}_{\mathrm{j}} \tag{27}$$

$$\frac{dR^{\mathrm{M}}_{9}}{dt} = \left(1 - F^{\mathrm{RdRp}}_{\mathrm{M}}\right)\frac{k^{\mathrm{Syn}}_{\mathrm{M}}Vp^{\mathrm{Nuc}}_{9}}{L_{9}} - k^{\mathrm{Deg}}_{\mathrm{M}}R^{\mathrm{M}}_{9} \tag{28}$$

with j = 1,...,8

$$Vp^{\mathrm{Nuc}}_{2} = Vp^{\mathrm{Nuc}}_{3} = Vp^{\mathrm{Nuc}}_{4} = Vp^{\mathrm{Nuc}}_{5} = Vp^{\mathrm{Nuc}}_{6} = Vp^{\mathrm{Nuc}}_{8} \tag{29}$$

$$f_{\mathrm{M}} = \begin{cases} F^{\mathrm{RdRp}}_{\mathrm{M}}, & \mathrm{j} \in \{1,\ldots,3\}, \\ 1, & \mathrm{j} \in \{4,\ldots,8\}, \end{cases} \tag{30}$$

$$k^{\mathrm{Syn}}_{\mathrm{M}} = \begin{cases} k^{\mathrm{Syn}}_{\mathrm{M,prim}}, & P_{\mathrm{RdRp}} < 1, \\ k^{\mathrm{Syn}}_{\mathrm{M,full}}, & P_{\mathrm{RdRp}} \geq 1, \end{cases} \tag{31}$$

$$\frac{dP_{\mathrm{PB1}}}{dt} = \frac{k_{\mathrm{P}}^{\mathrm{Syn}}}{D_{\mathrm{Rib}}} R_2^{\mathrm{M}} - k^{\mathrm{RdRp}} P_{\mathrm{PB1}} P_{\mathrm{PB2}} P_{\mathrm{PA}} \tag{32}$$

$$\frac{dP_{\mathrm{PB2}}}{dt} = \frac{k_{\mathrm{P}}^{\mathrm{Syn}}}{D_{\mathrm{Rib}}} R_1^{\mathrm{M}} - k^{\mathrm{RdRp}} P_{\mathrm{PB1}} P_{\mathrm{PB2}} P_{\mathrm{PA}} \tag{33}$$

$$\frac{dP_{\mathrm{PA}}}{dt} = \frac{k_{\mathrm{P}}^{\mathrm{Syn}}}{D_{\mathrm{Rib}}} R_3^{\mathrm{M}} - k^{\mathrm{RdRp}} P_{\mathrm{PB1}} P_{\mathrm{PB2}} P_{\mathrm{PA}} \tag{34}$$

$$\frac{dP_{\mathrm{RdRp}}}{dt} = k^{\mathrm{RdRp}} P_{\mathrm{PB1}} P_{\mathrm{PB2}} P_{\mathrm{PA}} - k_{\mathrm{V}}^{\mathrm{Syn}} P_{\mathrm{RdRp}} \sum_{\mathrm{i}} (Cp_{\mathrm{i}}) - k_{\mathrm{C}}^{\mathrm{Syn}} P_{\mathrm{RdRp}} \sum_{\mathrm{i}} \left(Vp_{\mathrm{i}}^{\mathrm{Nuc}}\right) \tag{35}$$

$$\frac{dP_{\mathrm{NP}}}{dt} = \frac{k_{\mathrm{P}}^{\mathrm{Syn}}}{D_{\mathrm{Rib}}} R_5^{\mathrm{M}} - \frac{k_{\mathrm{NP}}^{\mathrm{Bind}} P_{\mathrm{NP}}}{N_{\mathrm{NP}}^{\mathrm{Nuc}}} \sum_{\mathrm{i}} L_{\mathrm{V,i}} \left(R_{\mathrm{RdRp,i}}^{\mathrm{V}} + R_{\mathrm{RdRp,i}}^{\mathrm{C}}\right) \tag{36}$$

$$\begin{aligned} \frac{dP_{\mathrm{M1\text{-}STV}}}{dt} = {} & \frac{k_{\mathrm{P}}^{\mathrm{Syn}}}{D_{\mathrm{Rib}}} (1 - F_{\mathrm{Spl7}}) R_7^{\mathrm{M}} - \frac{k_{\mathrm{M1}}^{\mathrm{Bind}} P_{\mathrm{M1\text{-}STV}}}{N_{\mathrm{M1}}^{\mathrm{Nuc}}} \sum_{\mathrm{i}} \left(L_{\mathrm{V,i}} Vp_{\mathrm{i}}^{\mathrm{Nuc}}\right) - \frac{P_{\mathrm{M1\text{-}STV}}}{P_{\mathrm{M1,Tot}}} \left[\left(N_{P_{\mathrm{M1}}} - \frac{1}{N_{\mathrm{M1}}^{\mathrm{Nuc}}} \sum_{\mathrm{j}} L_{\mathrm{V,j}}\right) r_{\mathrm{STV}}^{\mathrm{Rel}} \right. \\ & \left. - \left(N_{P_{\mathrm{M1}}} - \frac{1}{N_{\mathrm{M1}}^{\mathrm{Nuc}}} \sum_{\mathrm{g}=1,\ldots,6,8,9} L_{\mathrm{V,g}}\right) r_{\mathrm{DIP}}^{\mathrm{Rel}} - \left(N_{P_{\mathrm{M1}}} - \frac{L_{\mathrm{V,9}}}{N_{\mathrm{M1}}^{\mathrm{Nuc}}}\right) r_{\mathrm{OP7}}^{\mathrm{Rel}} \right] \end{aligned} \tag{37}$$

$$\begin{aligned} \frac{dP_{\mathrm{M1\text{-}OP7}}}{dt} = {} & \frac{k_{\mathrm{P}}^{\mathrm{Syn}}}{D_{\mathrm{Rib}}} (1 - F_{\mathrm{Spl7}}) R_9^{\mathrm{M}} - \frac{P_{\mathrm{M1\text{-}OP7}}}{P_{\mathrm{M1,Tot}}} \left[\left(N_{P_{\mathrm{M1}}} - \frac{1}{N_{\mathrm{M1}}^{\mathrm{Nuc}}} \sum_{\mathrm{j}} L_{\mathrm{V,j}}\right) r_{\mathrm{STV}}^{\mathrm{Rel}} \right. \\ & \left. - \left(N_{P_{\mathrm{M1}}} - \frac{1}{N_{\mathrm{M1}}^{\mathrm{Nuc}}} \sum_{\mathrm{g}=1,\ldots,6,8,9} L_{\mathrm{V,g}}\right) r_{\mathrm{DIP}}^{\mathrm{Rel}} - \left(N_{P_{\mathrm{M1}}} - \frac{L_{\mathrm{V,9}}}{N_{\mathrm{M1}}^{\mathrm{Nuc}}}\right) r_{\mathrm{OP7}}^{\mathrm{Rel}} \right] \end{aligned} \tag{38}$$

$$\text{with } P_{\mathrm{M1,Tot}} = P_{\mathrm{M1\text{-}STV}} + P_{\mathrm{M1\text{-}OP7}} \tag{39}$$

$$\frac{dP_{\mathrm{NEP}}}{dt} = \frac{k_{\mathrm{P}}^{\mathrm{Syn}}}{D_{\mathrm{Rib}}} F_{\mathrm{Spl8}} R_8^{\mathrm{M}} - k^{\mathrm{Exp}} P_{\mathrm{NEP}} \sum_{\mathrm{i}} Vp_{\mathrm{M1,i}}^{\mathrm{Nuc}} \tag{40}$$

$$\frac{dP_{\mathrm{HA}}}{dt} = \frac{k_{\mathrm{P}}^{\mathrm{Syn}}}{D_{\mathrm{Rib}}} R_4^{\mathrm{M}} - N_{P_{\mathrm{HA}}} \left(r_{\mathrm{STV}}^{\mathrm{Rel}} + r_{\mathrm{DIP}}^{\mathrm{Rel}} + \frac{r_{\mathrm{OP7}}^{\mathrm{Rel}}}{N_{\mathrm{OP7}}^{\mathrm{Par}}}\right) \tag{41}$$

$$\frac{dP_{\mathrm{NA}}}{dt} = \frac{k_{\mathrm{P}}^{\mathrm{Syn}}}{D_{\mathrm{Rib}}} R_6^{\mathrm{M}} - N_{P_{\mathrm{NA}}} \left(r_{\mathrm{STV}}^{\mathrm{Rel}} + r_{\mathrm{DIP}}^{\mathrm{Rel}} + \frac{r_{\mathrm{OP7}}^{\mathrm{Rel}}}{N_{\mathrm{OP7}}^{\mathrm{Par}}}\right) \tag{42}$$

$$\frac{dP_{\mathrm{M2}}}{dt} = \frac{k_{\mathrm{P}}^{\mathrm{Syn}}}{D_{\mathrm{Rib}}} F_{\mathrm{Spl7}} \left(R_7^{\mathrm{M}} + R_9^{\mathrm{M}}\right) - N_{P_{\mathrm{M2}}} \left(r_{\mathrm{STV}}^{\mathrm{Rel}} + r_{\mathrm{DIP}}^{\mathrm{Rel}} + \frac{r_{\mathrm{OP7}}^{\mathrm{Rel}}}{N_{\mathrm{OP7}}^{\mathrm{Par}}}\right) \tag{43}$$

<u>Viral transcription and protein synthesis:</u> Nuclear vRNPs are also used as templates for the transcription of viral messenger RNA (mRNA) $R_{\mathrm{i}}^{\mathrm{M}}$ and each of the genome segments encodes for a different mRNA. The transcription of mRNAs depends on their respective length ($L_{\mathrm{i}}$) and is reduced for segments encoding for RdRp-related proteins, i.e., STV

segments 1 to 3, by the factor $F_{\mathrm{M}}^{\mathrm{RdRp}}$. Additionally, the generation of S7-OP7 mRNA is reduced by the factor $F_{\mathrm{M}}^{\mathrm{OP7}}$ to take into account the impact of the single-nucleotide substitutions (SNSs) on OP7 transcription. The availability of free viral polymerase $P_{\mathrm{RdRp}}$ is used as an indicator if mRNA transcription is occurring at the speed of primary transcription $\left(k_{\mathrm{M,prim}}^{\mathrm{Syn}}\right)$ or at full speed $\left(k_{\mathrm{M,full}}^{\mathrm{Syn}}\right)$. mRNAs are degraded with the rate $k_{\mathrm{M}}^{\mathrm{Deg}}$ and translated into viral proteins $P_{\mathrm{i}}$ in the cytoplasm. The polymerase sub-unit proteins ($P_{\mathrm{PB1}}$, $P_{\mathrm{PB2}}$, and $P_{\mathrm{PA}}$) bind to form RdRp. The mRNA of segment 7 can be translated into two different proteins, i.e., M1 and M2, and their production ratio is defined by $F_{\mathrm{Spl7}}$. S7-STV and S7-OP7 are translated into two different M1 proteins ($P_{\mathrm{M1\text{-}STV}}$ and $P_{\mathrm{M1\text{-}OP7}}$), and only M1-STV can bind vRNPs in the nucleus. However, we assume both S7-STV and S7-OP7 can be translated into regular M2. The proteins M1-STV, M1-OP7, M2, hemagglutinin (HA) and neuraminidase (NA) are required for virion release as they perform structural functions in progeny virions.

$$\frac{dV_{\mathrm{Cplx}}^{\mathrm{Cyt}}}{dt} = k^{\mathrm{Cplx}} Vp_{\mathrm{M1,7}}^{\mathrm{Cyt}} Vp_{\mathrm{M1,1}}^{\mathrm{Cyt}} 6Vp_{\mathrm{M1,5}}^{\mathrm{Cyt}} - r_{\mathrm{STV}}^{\mathrm{Rel}} - k_{\mathrm{Rnp}}^{\mathrm{Deg}} V_{\mathrm{Cplx}}^{\mathrm{Cyt}} \tag{44}$$

$$\frac{dD_{\mathrm{Cplx}}^{\mathrm{Cyt}}}{dt} = k^{\mathrm{Cplx}} Vp_{\mathrm{M1,9}}^{\mathrm{Cyt}} Vp_{\mathrm{M1,1}}^{\mathrm{Cyt}} 6Vp_{\mathrm{M1,5}}^{\mathrm{Cyt}} - r_{\mathrm{DIP}}^{\mathrm{Rel}} - k_{\mathrm{Rnp}}^{\mathrm{Deg}} D_{\mathrm{Cplx}}^{\mathrm{Cyt}} \tag{45}$$

$$\frac{dV^{\mathrm{Rel}}}{dt} = r_{\mathrm{STV}}^{\mathrm{Rel}} = k^{\mathrm{Rel}} \frac{V_{\mathrm{Cplx}}^{\mathrm{Cyt}}}{V_{\mathrm{Cplx}}^{\mathrm{Cyt}} + D_{\mathrm{Cplx}}^{\mathrm{Cyt}} + Vp_{\mathrm{M1,9}}^{\mathrm{Cyt}} + K_{V^{\mathrm{Rel}}}} \prod_{\mathrm{p}} \frac{P_{\mathrm{p}}}{P_{\mathrm{p}} + N_{P_{\mathrm{p}}} K_{V^{\mathrm{Rel}}}} \tag{46}$$

$$\frac{dD^{\mathrm{Rel}}}{dt} = r_{\mathrm{DIP}}^{\mathrm{Rel}} = k^{\mathrm{Rel}} \frac{D_{\mathrm{Cplx}}^{\mathrm{Cyt}}}{V_{\mathrm{Cplx}}^{\mathrm{Cyt}} + D_{\mathrm{Cplx}}^{\mathrm{Cyt}} + Vp_{\mathrm{M1,9}}^{\mathrm{Cyt}} + K_{V^{\mathrm{Rel}}}} \prod_{\mathrm{p}} \frac{P_{\mathrm{p}}}{P_{\mathrm{p}} + N_{P_{\mathrm{p}}} K_{V^{\mathrm{Rel}}}} \tag{47}$$

$$\frac{dOP7^{\mathrm{Rel}}}{dt} = r_{\mathrm{OP7}}^{\mathrm{Rel}} = \frac{k^{\mathrm{Rel}}}{N_{\mathrm{OP7}}^{\mathrm{Par}}} \frac{Vp_{\mathrm{M1,9}}^{\mathrm{Cyt}}}{V_{\mathrm{Cplx}}^{\mathrm{Cyt}} + D_{\mathrm{Cplx}}^{\mathrm{Cyt}} + Vp_{\mathrm{M1,9}}^{\mathrm{Cyt}} + K_{V^{\mathrm{Rel}}}} \prod_{\mathrm{p}} \frac{P_{\mathrm{p}}}{P_{\mathrm{p}} + N_{P_{\mathrm{p}}} K_{V^{\mathrm{Rel}}}} \tag{48}$$

$$\text{with } \mathrm{p} \in \{\mathrm{HA,NA,M1,M2}\}$$

$$\frac{dV_{\mathrm{Inf}}^{\mathrm{Rel}}}{dt} = F_{\mathrm{Par}} r_{\mathrm{STV}}^{\mathrm{Rel}} \tag{49}$$

$$\frac{dOP7_{\mathrm{Inf}}^{\mathrm{Rel}}}{dt} = F_{\mathrm{Par}} r_{\mathrm{OP7}}^{\mathrm{Rel}} \tag{50}$$

$$\frac{dV_{\mathrm{Tot}}^{\mathrm{Rel}}}{dt} = r_{\mathrm{STV}}^{\mathrm{Rel}} + r_{\mathrm{DIP}}^{\mathrm{Rel}} + r_{\mathrm{OP7}}^{\mathrm{Rel}} \tag{51}$$

Complex formation and virus particle release: The packaging of vRNPs in the cytoplasm is represented by a segment-specific mechanism. Cytoplasmic vRNPs form complexes including one copy of each genome segment. STV and "7+1" DIP complexes ($V_{\mathrm{Cplx}}^{\mathrm{Cyt}}$, $D_{\mathrm{Cplx}}^{\mathrm{Cyt}}$) both contain STV S1-6 and S8. In addition, they carry either a S7-STV or a S7-OP7 vRNP.

Complex formation occurs with rate $k^{\mathrm{Cplx}}$ and degradation with rate $k_{\mathrm{Rnp}}^{\mathrm{Deg}}$. In total, we consider the release of three different virus particles, i.e., STV particles, “7+1” DIPs and particles that only contain S7-OP7. Their release is described by the rates $r_{\mathrm{STV}}^{\mathrm{Rel}}$, $r_{\mathrm{DIP}}^{\mathrm{Rel}}$ and $r_{\mathrm{OP7}}^{\mathrm{Rel}}$, respectively. These rates are calculated considering the maximum release rate $k^{\mathrm{Rel}}$, the available vRNP complexes and the abundance of viral proteins. To calculate the absolute amount of released OP7 particles $r_{\mathrm{OP7}}^{\mathrm{Rel}}$, the released amount of S7-OP7 vRNPs is divided by the amount of S7-OP7 vRNPs included in one OP7 particle, i.e., $N_{\mathrm{OP7}}^{\mathrm{Par}}$. Here, we assume that $N_{\mathrm{OP7}}^{\mathrm{Par}} = 8$ molecules, analogous to the number of genome segments in a STV particle. Furthermore, we use $V_{\mathrm{Inf}}^{\mathrm{Rel}}$, $OP7_{\mathrm{Inf}}^{\mathrm{Rel}}$ and $V_{\mathrm{Tot}}^{\mathrm{Rel}}$ to represent the generated amount of infectious STV and OP7 particles as well as the total amount of released virions, respectively. The percentage of fully functional STV and OP7 is determined by the parameter $F_{\mathrm{Par}}$, which assumes that only a fraction of produced progeny particles is infectious (10).

simulation of intracellular model only:

$$\frac{dG_{\mathrm{IFN}}}{dt} = \frac{k_{\mathrm{Detec}}}{1 + \frac{P_{\mathrm{NS1}}}{K_{\mathrm{P}}}} \left(V^{\mathrm{Cyt}} + D^{\mathrm{Cyt}}\right) + k_{\mathrm{I,in}}^{\mathrm{Syn}} G_{\mathrm{JAK}} - k_{\mathrm{I}}^{\mathrm{Deg}} G_{\mathrm{IFN}} \tag{52.1}$$

simulation of multiscale model:

$$\frac{dG_{\mathrm{IFN}}}{dt} = -k_{\mathrm{I}}^{\mathrm{Deg}} G_{\mathrm{IFN}} \tag{52.2}$$

$$\frac{dG_{\mathrm{JAK}}}{dt} = \frac{k_{\mathrm{J}}^{\mathrm{Syn}}}{1 + e^{t-\tau_{\mathrm{J}}}} G_{\mathrm{IFN}} - k_{\mathrm{J}}^{\mathrm{Deg}} G_{\mathrm{JAK}} \tag{53}$$

$$\frac{dG_{\mathrm{MxA}}}{dt} = k_{\mathrm{MxA}}^{\mathrm{Syn}} G_{\mathrm{JAK}} - k_{\mathrm{MxA}}^{\mathrm{Deg}} G_{\mathrm{MxA}} \tag{54}$$

Interferon system: The interferon (IFN) system is implemented by considering three key components, i.e., IFN-β ($G_{\mathrm{IFN}}$), JAK/STAT ($G_{\mathrm{JAK}}$) and MxA ($G_{\mathrm{MxA}}$). The synthesis and degradation of these components is described by the rates $k_{\mathrm{I,in}}^{\mathrm{Syn}}$, $k_{\mathrm{J}}^{\mathrm{Syn}}$, $k_{\mathrm{MxA}}^{\mathrm{Syn}}$ and $k_{\mathrm{I}}^{\mathrm{Deg}}$, $k_{\mathrm{J}}^{\mathrm{Deg}}$, $k_{MxA}^{\mathrm{Deg}}$, respectively. The detection of incoming viral genomes by pattern recognition receptors in represented by the rate $k_{\mathrm{Detec}}$. This process is inhibited by the viral NS1 protein $P_{\mathrm{NS1}}$ considering $K_{\mathrm{P}}$, i.e., the amount of MxA required to reduce the rate of detection by half. Produced IFN-β is secreted and can attach to IFN-α/β receptors of the releasing cells (autocrine) and neighboring cells (paracrine). This induces the recruitment and expression of JAK/STAT, which initiates further IFN-β and MxA synthesis. We implemented a temporal feedback loop that reduces JAK/STAT production after a time $\tau_{\mathrm{J}}$, which is downregulated following initial activation (22-24). When simulating only the intracellular model for the initial parameter fit, IFN-β synthesis occurs as described in Equation (52.1) with an initial condition of $G_{\mathrm{IFN}}(t = 0) = 0$. When simulating the full multiscale model, Equation (52.2) is used with an initial condition of $G_{\mathrm{IFN}}(t = 0) = IFN(\tau)$ taking the current IFN-β value at time $\tau$ from the cell population level (see Equation (83)). In this case, we disregard the detection of viral genomes on the intracellular level, but consider the activation of the IFN system on the cell population level, which can lead to the protection of uninfected cells from productive infection.

**Cell population level**

$$\frac{dT}{dt} = \left(\mu - r_{\mathrm{STV}}^{\mathrm{Inf}} - r_{\mathrm{DIP}}^{\mathrm{Inf}} - k_{\mathrm{T}}^{\mathrm{Apo}}\right)T \tag{55}$$

$$\frac{dI_{\mathrm{DIP}}}{dt} = r_{\mathrm{DIP}}^{\mathrm{Inf}}T + \left(\mu - r_{\mathrm{STV}}^{\mathrm{Inf}} - k_{\mathrm{T}}^{\mathrm{Apo}}\right)I_{\mathrm{DIP}} \tag{56}$$

$$\text{with } \mu(t) = \left(\frac{\mu_{\mathrm{Max}}}{T_{\mathrm{Max}}}\left[T_{\mathrm{Max}} - C_{\mathrm{Tot}}(t)\right]\right)_{+} \tag{57}$$

$$\frac{\partial I_{\mathrm{STV}}}{\partial t} + \frac{\partial I_{\mathrm{STV}}}{\partial \tau} = -\left[r_{\mathrm{DIP}}^{\mathrm{Inf}} + k_{\mathrm{I}}^{\mathrm{Apo}}(\tau)\right]I_{\mathrm{STV}}(t,\tau) \tag{58}$$

$$\frac{\partial I_{\mathrm{CO}}}{\partial t} + \frac{\partial I_{\mathrm{CO}}}{\partial \tau} = -k_{\mathrm{I}}^{\mathrm{Apo}}(\tau)I_{\mathrm{CO}}(t,\tau) \tag{59}$$

$$\text{with } k_{\mathrm{I}}^{\mathrm{Apo}}(t) = \frac{K_{\mathrm{I}}}{1 + \exp\left[-\nu_{\mathrm{Apo}}\left(\tau - \tau_{\mathrm{Apo}}\right)\right]} \tag{60}$$

$$\frac{dT_{\mathrm{A}}}{dt} = k_{\mathrm{T}}^{\mathrm{Apo}}T - \left(r_{\mathrm{STV}}^{\mathrm{Inf}} + r_{\mathrm{DIP}}^{\mathrm{Inf}} + k^{\mathrm{Lys}}\right)T_{\mathrm{A}} \tag{61}$$

$$\frac{dI_{\mathrm{A}}}{dt} = \int_0^\infty k_{\mathrm{I}}^{\mathrm{Apo}}(\tau)\left[I_{\mathrm{STV}}(t,\tau) + I_{\mathrm{CO}}(t,\tau)\right]d\tau + k_{\mathrm{T}}^{\mathrm{Apo}}I_{\mathrm{DIP}} + \left(r_{\mathrm{STV}}^{\mathrm{Inf}} + r_{\mathrm{DIP}}^{\mathrm{Inf}}\right)T_{\mathrm{A}} - k^{\mathrm{Lys}}I_{\mathrm{A}} \tag{62}$$

$$C_{\mathrm{Tot}}(\tau) = T(t) + T_{\mathrm{A}}(t) + I_{\mathrm{DIP}}(t) + \int_0^\infty I_{\mathrm{STV}}(t,\tau)\,d\tau + \int_0^\infty I_{\mathrm{CO}}(t,\tau)\,d\tau + I_{\mathrm{A}}(t) \tag{63}$$

Cell populations: On the cell population level, the model considers populations of uninfected cells ($T$), STV-only infected cells ($I_{\mathrm{STV}}$), OP7-only infected cells ($I_{\mathrm{DIP}}$), coinfected cells ($I_{\mathrm{CO}}$), apoptotic uninfected cells ($T_{\mathrm{A}}$), and apoptotic infected cells ($I_{\mathrm{A}}$). To facilitate comparability to previous versions of the multiscale model (15), we define OP7 particles as “DIP” on the population level and use this wording in the state and parameter nomenclature ($I_{\mathrm{DIP}}$, etc.). STV-only and coinfected cells are considered as age-segregated populations and are classified by their infection age $\tau$. Uninfected cells proliferate with the specific growth rate $\mu$, can get infected by STV and OP7 with rates $r_{\mathrm{STV}}^{\mathrm{Inf}}$ and $r_{\mathrm{DIP}}^{\mathrm{Inf}}$, respectively, and get apoptotic with rate $k_{\mathrm{T}}^{\mathrm{Apo}}$. Similarly, OP7-only infected cells can grow, get infected by STV and undergo apoptosis. STV-only infected cells do not proliferate, but can get infected by OP7 and get apoptotic with the infection-induced apoptosis rate $k_{\mathrm{I}}^{\mathrm{Apo}}(\tau)$. This rate depends on the infection age $\tau$ and utilizes a logistic function, which can approximate the cumulative density function of the normal distribution (25). Coinfected cells are also subject to infection-induced apoptosis. Apoptotic uninfected cells can become infected by either STV or OP7 and undergo cell lysis with rate $k^{\mathrm{Lys}}$. The same lysis rate applies to apoptotic infected cells. The total amount of viable cells is collected as $C_{\mathrm{Tot}}$ in Equation (62).

<u>Infection rates and superinfection exclusion</u>: The infection rates of cells are separated for STV and OP7, resulting in

$$r_{\mathrm{STV}}^{\mathrm{Inf}} = F_{\mathrm{Inf}} k^{\mathrm{Fus}} V^{\mathrm{En}} \theta\left(\frac{1}{T + T_{\mathrm{A}} + I_{\mathrm{DIP}}}\right) \tag{64}$$

$$r_{\mathrm{DIP}}^{\mathrm{Inf}} = F_{\mathrm{Inf}} k^{\mathrm{Fus}} D^{\mathrm{En}} \theta\left(\frac{1}{T + T_{\mathrm{A}} + \int_0^{\tau_{\mathrm{SIE}}} I_{\mathrm{STV}}(t,\tau) d\tau}\right) \tag{65}$$

$$\text{with } \theta(a) = \begin{cases} a, & a > 0, \\ 0, & a \leq 0, \end{cases} \tag{66}$$

where STV and OP7 in endosomes are denoted by $V^{\mathrm{En}}$ and $D^{\mathrm{En}}$, respectively. Their amount is related to the fusion rate $k^{\mathrm{Fus}}$ and the fraction of cells successfully infected by fusion of a single virion $F_{\mathrm{Inf}}$. To account for the current number of suitable infection targets, this is divided by the total amount of cells that are targeted by either STV or OP7. The parameter $\tau_{\mathrm{SIE}}$ describes the infection age after which STV-only infected cells cannot get re-infected by OP7.

Target cells $T$ and their apoptotic counterpart $T_{\mathrm{A}}$ can be infected by either STV or OP7, however, we assume re-infection of STV- and OP7-only infected cells ($I_{\mathrm{STV}}$ and $I_{\mathrm{DIP}}$) is only possible by the opposing virus particle. Additionally, STV-only infected cells are protected from re-infection by OP7 after a certain time period $\tau_{\mathrm{SIE}}$ to consider superinfection exclusion, which is mediated by neuraminidase (26). According to (27), we set the time window for a productive coinfection of STV-only infected cells to $\tau_{\mathrm{SIE}} = 3$ h after initial infection. This results in STV-only infected cells that have reached an infection age of 3 h to remain in this state and continuously producing progeny STV. This implementation prevents that OP7 is produced at later stages to transform all STV-only infected cells to coinfected cells, which would result in an inaccurate description of STV titers.

Furthermore, we assume that STV-only infected cells are not re-infected by STV, OP7-only infected cells are not re-infected by OP7, and coinfected cells are not re-infected by either virus particle. This was implemented to prevent the existence of a continuous sink draining virus particles from the system. Lastly, we assume that OP7-only infected cells can get re-infected by STV and converted to coinfected cells at all times, because they are not capable of replication as they cannot produce progeny RdRp. This would likely result in low levels of neuraminidase, which are not sufficient to prevent re-infection.

Age-segregated infected cell populations: For the model of STV and OP7 coinfection, we consider age-segregated populations for STV-only and coinfected cells.

$$I_{\mathrm{STV}}(t,\tau) = \begin{cases} r_{\mathrm{STV}}^{\mathrm{Inf}}(t-\tau)T(t-\tau)\exp\left[-\int_0^{\tau} r_{\mathrm{DIP}}^{\mathrm{Inf}}(a) + k_{\mathrm{I}}^{\mathrm{Apo}}(a)da\right], & t > \tau \geq 0, \\ 0, & \tau > t \geq 0, \end{cases} \quad (67)$$

$$I_{\mathrm{CO}}(t,\tau) = \begin{cases} \left[r_{\mathrm{STV}}^{\mathrm{Inf}}(t-\tau)I_{\mathrm{DIP}}(t-\tau) + r_{\mathrm{DIP}}^{\mathrm{Inf}}(t-\tau)\int_0^{\tau_{\mathrm{SIE}}} I_{\mathrm{STV}}(t-\tau,a)da\right]\exp\left[-\int_0^{\tau} k_{\mathrm{I}}^{\mathrm{Apo}}(a)da\right], & t > \tau \geq 0, \\ 0, & \tau > t \geq 0, \end{cases}$$

$$(68)$$

These algebraic equations can be derived from Equations (58) and (59) providing the infection age density for both species. Upon infection at time $t_{\mathrm{I}}$, corresponding cells with age zero emerge. This leads to boundary condition of $I_{\mathrm{STV}}(t_{\mathrm{I}},\tau=0) = r_{\mathrm{STV}}^{\mathrm{Inf}}(t_{\mathrm{I}})T(t_{\mathrm{I}})$ and $I_{\mathrm{CO}}(t_{\mathrm{I}},\tau=0) = r_{\mathrm{STV}}^{\mathrm{Inf}}(t_I)I_{\mathrm{DIP}}(t_{\mathrm{I}}) + r_{\mathrm{DIP}}^{\mathrm{Inf}}(t_{\mathrm{I}})I_{\mathrm{STV}}(t_{\mathrm{I}})$ for STV-only and coinfected cells, respectively. Both types of populations are subject to infection-induced apoptosis, however, STV-only infected cells can also still get infected by OP7.

Virus particle release: Infectious STV ($V$) and OP7 ($D$) can theoretically be released from both STV-only and coinfected cells. OP7-only infected cells are assumed to not release any virus particles. Therefore, we obtain the following release dynamics

$$\frac{dV}{dt} = \int_0^{\infty}\left[r_{\mathrm{STV},I_{\mathrm{STV}}}^{\mathrm{Rel}}(\tau)I_{\mathrm{STV}}(t,\tau) + r_{\mathrm{STV},I_{\mathrm{CO}}}^{\mathrm{Rel}}(\tau)I_{\mathrm{CO}}(t,\tau)\right]d\tau - k_{\mathrm{V}}^{\mathrm{Deg}}V + \sum_{\mathrm{n}}\left(k_{\mathrm{n}}^{\mathrm{Dis}}V_{\mathrm{Hi}}^{\mathrm{Att}} - k_{\mathrm{c,n}}^{\mathrm{Att}}B_{\mathrm{n}}^{\mathrm{V}}V\right) \quad (69)$$

$$\frac{dD}{dt} = \int_0^{\infty}\left[r_{\mathrm{OP7},I_{\mathrm{STV}}}^{\mathrm{Rel}}(\tau)I_{\mathrm{STV}}(t,\tau) + r_{\mathrm{OP7},I_{\mathrm{CO}}}^{\mathrm{Rel}}(\tau)I_{\mathrm{CO}}(t,\tau)\right]d\tau - k_{\mathrm{V}}^{\mathrm{Deg}}D + \sum_{\mathrm{n}}\left(k_{\mathrm{n}}^{\mathrm{Dis}}D_{\mathrm{Hi}}^{\mathrm{Att}} - k_{\mathrm{c,n}}^{\mathrm{Att}}B_{\mathrm{n}}^{\mathrm{D}}D\right) \quad (70)$$

$$\text{with } B_{\mathrm{n}}^{\mathrm{V}} = B_{\mathrm{n}}^{\mathrm{Tot}}(T + T_{\mathrm{A}} + I_{\mathrm{DIP}}) - V_{\mathrm{n}}^{\mathrm{Att}}, \quad (71)$$

$$B_{\mathrm{n}}^{\mathrm{D}} = B_{\mathrm{n}}^{\mathrm{Tot}}\left(T + T_{\mathrm{A}} + \int_0^{\tau_{\mathrm{SIE}}} I_{\mathrm{STV}}(t,\tau)d\tau\right) - D_{\mathrm{n}}^{\mathrm{Att}}, \quad (72)$$

$$\text{and } k_{\mathrm{n}}^{\mathrm{Dis}} = \frac{k_{\mathrm{c,n}}^{\mathrm{Att}}}{k_{\mathrm{c,n}}^{\mathrm{Eq}}}, \ \mathrm{n} \in \{\mathrm{Hi,Lo}\} \quad (73)$$

where STV and OP7 attach to their free binding sites on the cell membrane with rate $k_{\mathrm{c,n}}^{\mathrm{Att}}$. We assume STV does not re-infect STV-only and coinfected cells, while OP7 do not re-infect

OP7-only and coinfected cells. This limits the potential binding sites for the two different virus particles, i.e. $B_{\mathrm{n}}^{\mathrm{V}}$ and $B_{\mathrm{n}}^{\mathrm{D}}$, as described in Equations (71) and (72). Attached STV ($V_{\mathrm{n}}^{\mathrm{Att}}$) and OP7 ($D_{\mathrm{n}}^{\mathrm{Att}}$) can detach from binding sites with rate $k_{\mathrm{n}}^{\mathrm{Dis}}$ based on the equilibrium constant $k_{\mathrm{c,n}}^{\mathrm{Eq}}$. Additionally, they are degraded with rate $k_{\mathrm{V}}^{\mathrm{Deg}}$ which is applied to both STV and OP7. The rates $r_{\mathrm{i,j}}^{\mathrm{Rel}}(\tau)$ describe virus particle release of i ∈ {STV,DIP} particles from j ∈ $\{I_{\mathrm{STV}}, I_{\mathrm{CO}}\}$ cells.

For the description of experimental data, we also consider the total amounts of released STV ($V_{\mathrm{Tot}}$), OP7 ($D_{\mathrm{Tot}}$), and the total number of released virus particles ($P_{\mathrm{Tot}}$)

$$\frac{dV_{\mathrm{Tot}}}{dt} = \int_0^{\infty} \left[r_{\mathrm{STV},I_{\mathrm{STV}},\mathrm{Tot}}^{\mathrm{Rel}}(\tau) I_{\mathrm{STV}}(t,\tau) + r_{\mathrm{STV},I_{\mathrm{CO}},\mathrm{Tot}}^{\mathrm{Rel}}(\tau) I_{\mathrm{CO}}(t,\tau)\right] d\tau \quad (74)$$

$$\frac{dD_{\mathrm{Tot}}}{dt} = \int_0^{\infty} \left[r_{\mathrm{OP7},I_{\mathrm{STV}},\mathrm{Tot}}^{\mathrm{Rel}}(\tau) I_{\mathrm{STV}}(t,\tau) + r_{\mathrm{OP7},I_{\mathrm{CO}},\mathrm{Tot}}^{\mathrm{Rel}}(\tau) I_{\mathrm{CO}}(t,\tau)\right] d\tau \quad (75)$$

$$\frac{dP_{\mathrm{Tot}}}{dt} = \frac{dV_{\mathrm{Tot}}}{dt} + \frac{dD_{\mathrm{Tot}}}{dt} \quad (76)$$

<u>Virus entry</u>: Attached STV and OP7 undergo endocytosis with rate $k^{\mathrm{En}}$ and virus particles in endosomes fuse with the endosomal membrane with rate $k^{\mathrm{Fus}}$.

$$\frac{dV_{\mathrm{n}}^{\mathrm{Att}}}{dt} = k_{\mathrm{c,n}}^{\mathrm{Att}} B_{\mathrm{n}}^{\mathrm{V}} V - \left(k_{\mathrm{n}}^{\mathrm{Dis}} + k^{\mathrm{En}} + r_{\mathrm{STV}}^{\mathrm{Inf}} + r_{\mathrm{STV}}^{\mathrm{Lys}}\right) V_{\mathrm{n}}^{\mathrm{Att}} \quad (77)$$

$$\frac{dD_{\mathrm{n}}^{\mathrm{Att}}}{dt} = k_{\mathrm{c,n}}^{\mathrm{Att}} B_{\mathrm{n}}^{\mathrm{D}} D - \left(k_{\mathrm{n}}^{\mathrm{Dis}} + k^{\mathrm{En}} + r_{\mathrm{DIP}}^{\mathrm{Inf}} + r_{\mathrm{DIP}}^{\mathrm{Lys}}\right) D_{\mathrm{n}}^{\mathrm{Att}} \quad (78)$$

$$\frac{dV^{\mathrm{En}}}{dt} = k^{\mathrm{En}}\left(V_{\mathrm{Hi}}^{\mathrm{Att}} + V_{\mathrm{Lo}}^{\mathrm{Att}}\right) - \left(k^{\mathrm{Fus}} + r_{\mathrm{STV}}^{\mathrm{Inf}} + r_{\mathrm{STV}}^{\mathrm{Lys}}\right) V^{\mathrm{En}} \quad (79)$$

$$\frac{dD^{\mathrm{En}}}{dt} = k^{\mathrm{En}}\left(D_{\mathrm{Hi}}^{\mathrm{Att}} + D_{\mathrm{Lo}}^{\mathrm{Att}}\right) - \left(k^{\mathrm{Fus}} + r_{\mathrm{DIP}}^{\mathrm{Inf}} + r_{\mathrm{DIP}}^{\mathrm{Lys}}\right) D^{\mathrm{En}} \quad (80)$$

$$r_{\mathrm{STV}}^{\mathrm{Lys}} = k^{\mathrm{Lys}} T_{\mathrm{A}} \theta \left(\frac{1}{T + T_{\mathrm{A}} + I_{\mathrm{DIP}}}\right) \quad (81)$$

$$r_{\mathrm{DIP}}^{\mathrm{Lys}} = k^{\mathrm{Lys}} T_{\mathrm{A}} \theta \left(\frac{1}{T + T_{\mathrm{A}} + \int_0^{\infty} I_{\mathrm{STV}}(t,\tau) d\tau}\right) \quad (82)$$

During cell infection and lysis, virus particles that are attached to binding sites and inside of endosomes are also removed from the system according to the infection and lysis rates ($r_{\mathrm{STV}}^{\mathrm{Lys}}$ and $r_{\mathrm{DIP}}^{\mathrm{Lys}}$). The lysis rates are determined by relating the uninfected apoptotic cells undergoing lysis to the number of cells having STV or OP7 attached or in endosomes in Equations (81) and (82).

<u>Interferon system</u>: IFN-β is implemented on the extracellular levels assuming a constant

release from all living cells.

$$\frac{dIFN}{dt} = k_{\mathrm{I,base}}^{\mathrm{Syn}}T + \frac{k_{\mathrm{I,ex}}^{\mathrm{Syn}}}{(1 + e^{(t-\tau_{\mathrm{I}})})}\left(I_{\mathrm{STV}} + f_{\mathrm{IFN}}\left(I_{\mathrm{DIP}} + I_{\mathrm{CO}}\right)\right) - k_{\mathrm{IFN}}^{\mathrm{Deg}}IFN \quad (83)$$

$$f_{\mathrm{IFN}} = \begin{cases} F_{\mathrm{IFN}}^{\mathrm{high}}, & \mathrm{MODIP} \geq 500 \\ 1, & \mathrm{MODIP} < 500 \end{cases} \quad (84)$$

where $k_{\mathrm{I,base}}^{\mathrm{Syn}}$ and $k_{\mathrm{I,ex}}^{\mathrm{Syn}}$ describe rates of IFN-β release from uninfected and infected cells, respectively. $k_{\mathrm{I,base}}^{\mathrm{Syn}}$ is calibrated to establish a baseline level of IFN-β in the system when no cells are infected. $F_{\mathrm{IFN}}^{\mathrm{high}}$ defines the increased release of IFN-β from cells infected with a high OP7 dose. $\tau_{\mathrm{I}}$ represents the time after which IFN-β production is downregulated by feedback inhibition following infection (22-24). IFN-β degradation is denoted by the parameter $k_{\mathrm{IFN}}^{\mathrm{Deg}}$ (12).
